# Intranasal CRMP2-Ubc9 Inhibitor Regulates Na_V_1.7 to Alleviate Trigeminal Neuropathic Pain

**DOI:** 10.1101/2023.07.16.549195

**Authors:** Santiago I. Loya-Lopez, Heather N. Allen, Paz Duran, Aida Calderon-Rivera, Kimberly Gomez, Upasana Kumar, Rory Shields, Rui Zeng, Akshat Dwivedi, Saumya Saurabh, Olga A. Korczeniewska, Rajesh Khanna

## Abstract

Dysregulation of voltage-gated sodium Na_V_1.7 channels in sensory neurons contributes to chronic pain conditions, including trigeminal neuropathic pain. We previously reported that chronic pain results in part from increased SUMOylation of collapsin response mediator protein 2 (CRMP2), leading to an increased CRMP2/Na_V_1.7 interaction and increased functional activity of Na_V_1.7. Targeting this feed-forward regulation, we developed compound **194**, which inhibits CRMP2 SUMOylation mediated by the SUMO-conjugating enzyme Ubc9. We further demonstrated that **194** effectively reduces the functional activity of Na_V_1.7 channels in dorsal root ganglia neurons and alleviated inflammatory and neuropathic pain. Here, we employed a comprehensive array of investigative approaches, encompassing biochemical, pharmacological, genetic, electrophysiological, and behavioral analyses, to assess the functional implications of Na_V_1.7 regulation by CRMP2 in trigeminal ganglia (TG) neurons. We confirmed the expression of *Scn9a*, *Dpysl2*, and *UBE2I* within TG neurons. Furthermore, we found an interaction between CRMP2 and Na_V_1.7, with CRMP2 being SUMOylated in these sensory ganglia. Disrupting CRMP2 SUMOylation with compound **194** uncoupled the CRMP2/Na_V_1.7 interaction, impeded Na_V_1.7 diffusion on the plasma membrane, and subsequently diminished Na_V_1.7 activity. Compound **194** also led to a reduction in TG neuron excitability. Finally, when intranasally administered to rats with chronic constriction injury of the infraorbital nerve (CCI-ION), **194** significantly decreased nociceptive behaviors. Collectively, our findings underscore the critical role of CRMP2 in regulating Na_V_1.7 within TG neurons, emphasizing the importance of this indirect modulation in trigeminal neuropathic pain.

## 1. Introduction

The trigeminal ganglion (TG) innervates craniofacial areas via the ophthalmic, maxillary, and mandibular nerves [79] and plays a crucial role in craniofacial nociception [3]. Post-traumatic trigeminal pain (PTNP) is a severe unilateral pain with limited long-term improvement [63; 70] caused by dental procedures or craniofacial/oral traumas [31; 34; 40; 49; 51; 60; 64; 65; 71; 75]. Unfortunately, PTNP management drugs [29; 43; 80] have central side effects due to their broad molecular targets. Understanding the molecular components contributing to PTNP pathophysiology is crucial for developing more effective, targeted drugs.

Craniofacial noxious stimulation activates intracellular signaling cascades, altering membrane ionic channel activity [38; 73; 74], leading to increased peripheral nerve excitability. For instance, dysregulation of N-type voltage-gated calcium (Ca_V_2.2) channels has been linked to cephalic pain [23; 68]. Preclinical evidence suggests that inhibiting their interaction with collapsin response-mediated protein 2 (CRMP2) can reduce excitatory neurotransmitter release [62] and decrease periorbital allodynia [10; 54]. The T-type voltage-gated calcium (Ca_V_3.2) channels are also important mediators of trigeminal neuropathic pain [30; 59]. Patients with trigeminal neuralgia exhibit point mutations in the Ca_V_3.2 channel gene [19] leading to altered funtion of the channel [30; 59].Voltage-gated sodium channels (VGSCs) also play a critical role in trigeminal pain transduction [81; 82]. In rats with temporomandibular joint inflammation, Na_V_1.7 mRNA and protein levels increase in the TG [81], correlating with enhanced sodium currents in TG neurons [82]. However, no link between CRMP2 and Na_V_1.7 in the trigeminal system has been established.

We previously demonstrated CRMP2 regulation of Na_V_1.7 in dorsal root ganglion (DRG) neurons [14; 15; 20; 22; 26], with pain increasing CRMP2 expression [52; 53]. SUMOylation of CRMP2 enhances functional expression of Na_V_1.7 [20; 22], promoting DRG excitability [52; 53]. We have developed a small molecule called **194** that targets the CRMP2-Na_V_1.7 interaction [21; 27], and inhibits CRMP2 SUMOylation. By reducing trafficking and hyperactivity of Na_V_1.7 in DRG neurons across multiple species, including humans, **194** provides a non-opioid alternative for neuropathic pain treatment [9; 13; 44]. Our preclinical studies have shown that 194 decreases spinal nociceptive transmission, normalizing the response to mechanical stimulation in acute and neuropathic pain models in rodents [9; 13; 44].

Evidence points to the potential of investigating the CRMP2-Na_V_1.7 interaction in TG neurons to address trigeminal pain treatment. Based on shared characteristics between DRG and TG, our study explored if regulation of Na_V_1.7 by CRMP2 also occurs in TG neurons and if **194** can disrupt this regulation to alleviate PTNP. Our work revealed the presence and interaction of both CRMP2 and Na_V_1.7 in TGs and showed that exposing TG neurons to **194** decreased *(i)* difussion and surface expression of Na_V_1.7 channels, *(ii)* density of Na_V_1.7 currents, and *(ii)* TG neuron exctibaility. Lastly, we observed that intranasal administration of compound **194** reduced chronic constriction injury of the infraorbital nerve (ION-CCI)-induced nociceptive behaviors. These findings highlight the importance of the Na_V_1.7/CRMP2 interaction in TG sensory neurons and demonstrate the efficacy of **194** in inhibiting established neuropathic trigeminal pain.

## 2. Materials and methods

### 2.1 Animals

Pathogen-free adult male and female Sprague-Dawley rats (150-200 g; Charles River) were housed in temperature-controlled (23 ± 3 ⁰C) and light-controlled (12-h light/12-h dark cycle; lights on 7:00-19:00) rooms with standard rodent chow and water available *ad libitum.* The Institutional Animal Care and Use Committee of New York University approved all experiments. All procedures were conducted in accordance with the *Guide for Care and Use of Laboratory Animals* published by the National Institutes of Health.

For the surgical procedures and behavioral experiments: adult (56 days old, 362.55 ± 16.06 g, Charles River Laboratories) male and female Sprague Dawley rats were used. Rats were pair-housed in animal care system IVC M.I.C.E. cages with access to food and hyper-chlorinated reverse osmosis water *ad libitum*. Facility was maintained on a 12:12 light: dark cycle at 23 ± 3 ⁰C (humidity between 30-50%). Two wooden blocks and an enviro-pack was provided per cage for environmental enrichment. Experimental procedures were approved by the Rutgers University Institutional Animal Care and Use Committee protocol NO. 999901099. Animal experiments were performed in accordance with the International Association for the Study of Pain guidelines for research involving animals.

### 2.2 Chemicals

All chemicals, unless noted, were purchased from Sigma (St Louis, MO). **194** (benzoylated 2-(4-piperidinyl)-1,3-benzimidazole analog; molecular weight 567.6) was obtained from TCG Lifesciences (Kolkata, India) and >99% purity was confirmed using HPLC and resuspended in DMSO for in vitro use. The synthesis of **194** has been previously reported by us [13].

### 2.3 Isolation and culture of rat trigeminal ganglion (TG) neurons

Female Sprague-Dawley rats were euthanized through an isoflurane overdose and decapitated. Trigeminal ganglia were identified, collected, enzymatically digested in DMEM F12 media (cat. no. 11765, GIBCO) with dispase type II (2 mg mL −1, MB, cat. no. 165859) and collagenase type II (3 mg mL −1, cat. no. 4176, Worthington) for ∼40 min at 37 ⁰C under gentle agitation and centrifuged at 800 x g for 7 minutes. The supernatant was discarded, and the pellet was resuspended in complete TG media (DMEM F12 media containing 1% penicillin/streptomycin sulfate (10,000 µL^-1^, stock) and 10% fetal bovine serum (Hyclone)).

For the corresponding set of experiments, trigeminal ganglion neurons (TGs) were transfected with EGFP and Invitrogen Stealth RNAi™ siRNA Negative Control (scramble siRNA) (cat. no. 12935300, Thermo Fisher Scientific) or CRMP2 siRNA (5′ GTAAACTCCTTCCTCGTGT-3′; obtained from Thermo Fisher Scientific) using the 4D-Nucleofector (P3 Primary Cell Solution, program DR 114; Lonza Biosciences). We have previously validated efficiency of knockdown of CRMP2 protein by CRMP2 siRNA in a variety of neurons [12; 16; 56; 78]. For the total internal reflection (TIR) microscopy, TG trigeminal ganglion neurons were transfected with 3 µg of Na_V_1.7-PEPCy3 plasmid and 0.75 µg green fluorescent protein (GFP) using the rat neuron 4D-Nucleofector solution (P3 Primary Cell Solution, program DR 114; Lonza Biosciences). Neurons were plated on round bottom glass dishes (cat. no. P35G-1.5-10-C, Mattek) and maintained at 37 °C and 5% CO_2_ in complete DMEM F12 media. Successfully transfected cells were identified by GFP fluorescence.

### 2.4 *In situ* hybridization

TG tissue was extracted fresh and flash frozen on dry ice before being embedded in optimal cutting temperature gel (Cat. No. 23-730-571, Fisher Scientific). Tissue was sectioned at 20 µm on a cryostat at - 20°C and mounted directly onto SuperFrost Plus microscope slides (cat. no22-037-246, Fisher Scientific). Slides were air dried at room temperature for 24 hours before undergoing RNAScope *in situ* hybridization. Slides were immersed in distilled water for 5 minutes to remove OCT before immersion in chilled 10% neutral buffered formalin (STL286001, Fisher Scientific) for 15 minutes.

Slides then underwent dehydration in a series ethanol baths at 50% (5 min), 70% (5 min), and finally 100% (10 min) ethanol [32]. Slides were air dried, and a hydrophobic barrier was drawn around the tissue sections. RNAScope was carried out according to ACD instructions. Briefly, tissue was digested using Protease IV for 3 minutes at room temperature. Slides were washed in RNAScope wash buffer for 4 minutes before being incubated in RNAScope probes for 2 hours at 40C in HybEZ oven. Probes for *Rbfox3* (ACD Bio. 436351), *Scn9a* (ACD Bio. 317851), *Calca* (ACD Bio. 317511), *Nefh* (ACD Bio. 474241), *Dpysl2* (ACD Bio. 1235621), and *Ube2i* (ACD Bio. 1253901) were purchased from ACD Bio. Slides then underwent a series of amplification via incubation in AMP1 (30 min), AMP2 (30 min), and AMP3 (15 min) at 40°C, with 4-minute wash buffer baths at room temperature between each step. Each channel was developed for 15 minutes at 40°C (HRP C1, C2 or C3) and underwent a 4-minute bath in wash buffer at room temperature before incubation with a tyrosine signal amplification (TSA) based fluorophore for 30 minutes at 40°C. After a 4-minute wash buffer bath, HRP blocker was applied for 15 minutes at 40°C before developing the next channel. Slides were incubated with DAPI for 30 seconds before coverslipping with EverBrite (Biotium, 23003) hardset mounting medium and drying overnight at room temperature [32]. Stitched images encompassing the TG were collected on a Leica DMI8 microscope (Wetzlar, Germany) using a 40x objective and analyzed using QuPath software v0.4.3.

Only neurons were counted, defined by a DAPI-labeled nucleus surrounded by *Rbfox3* signal. Sub analysis quantified neurons expressing *Scn9a*, *Calca*, *Nefh*, *Dpysl2*, or *Ube2i*. Neurons were considered positive for a marker if they expressed greater than 10 fluorescent puncta. Each individual data point represents the mean of 4-6 sections from one animal.

### 2.5 Immunofluorescence, confocal microscopy, and quantification of Na_V_1.7

Female rat TG neurons in culture were incubated overnight with either DMSO (as control) or 5 µM of **194**. Immunofluorescence was performed as described previously [20; 28; 52; 58]. Briefly, cells were fixed for 5 min using ice-cold methanol and allowed to dry at room temperature. Cells were rehydrated in phosphate buffered saline (PBS) and incubated overnight with anti-Na_V_1.7 antibody (1/200; Cat #: MABN41, Sigma-Aldrich) in PBS with 3% bovine serum albumin at 4°C. Cells were then washed 3 times in PBS and incubated with PBS containing 3% bovine serum albumin and secondary antibody Alexa 594 goat anti-mouse (Thermo Fisher Scientific, Waltham, MA) for 1 hour at room temperature. After washing with PBS, cells were stained with 49,6-diamidino-2-phenylindole (DAPI, 50 mg/mL) and mounted in ProLong Diamond Antifade Mountant (Cat #: P36961, Life Technologies Corporation). Immunofluorescent micrographs were acquired using a Plan-Apochromat 63x/1.4 oil CS2 objective on a Leica SP8 confocal microscope operated by the LAS X microscope software (Leica). Camera gain and other relevant settings were kept constant. Membrane immunoreactivity was calculated by measuring the signal intensity in the area contiguous to the boundary of the cell. The membrane to cytosol ratio was determined by defining regions of interest in the cytosol and on the membrane of each cell using Image J. Total fluorescence was normalized to the area analyzed and before calculating the ratios. The experimenter was blinded to the experimental conditions.

### 2.6 Total internal reflection (TIR) microscopy of TG neurons

#### 2.6.1. Fusion of Na_V_1.7 to a Cy3 binding polypeptide PEPCy

Genetically encoded photostable dyes are a key requirement to observe single Na_V_1.7 dynamics for extended time scales. In this regard, we isolated single chain variable fragments (scFv) that bind the Cyanine dye Cy3 with <100 nM affinity. Briefly, a library of yeast cells optimized for surface expression of proteins was created by mutagenizing previously published dye [35] and fluorogen binding scFvs [66; 72]. Cy3 binding clones were selected via flow cytometry, enriched, and further mutagenized to obtain the 29 kDa scFv refered to as Photostability Enhancing Peptide (PEPCy) in this study. To create the Na_V_1.7-PEPCy3 plasmid, the complete open reading frame of mouse Na_V_1.7 was cloned into the pcDNA3.1(+)-C-DYK vector (GenScript) with several modifications. The PEPCy3 tag was fused in-frame at the N-terminus of the Na_V_1.7 sequence using a small 8 amino acid G linker. Additionally, to promote membrane localization and orient the PEPCy3 toward the extracellular membrane, the N-terminus portion of β subunit was fused to the N-terminus of the full sequence.

#### 2.6.2. Total internal reflection (TIR) microscopy of TG neurons

TG neurons transfected with the Na_V_1.7-PEPCy scFv plasmid were plated on round bottom glass dishes (cat. no. P35G-1.5-10-C, Mattek) were incubated with 0.1% DMSO or compound **194** (5 µM) overnight at 37°C. Prior to imaging, the cells were washed 1x with 1mL Colorless Tyrode’s buffer followed by incubating the samples with 50 nM Sulfo-Cy3 carboxylic acid (Lumiprobe) for 20 mins at 37°C. The cells were gently washed 2x with colorless Tyrode solution followed by imaging in the same medium with or without compound **194** as appropriate. The mattek dish containing TG neurons was then placed in a temperature and humidity-controlled chamber (OkoLabs) on a custom-built microscope stand (Nikon). Microscopy was performed at 37°C. An excitation laser line of 561 nm was used with approximate intensity of 500 W/cm^2^ at the sample. For total internal reflection (TIR) microscopy, a super apochromat objective (PlanApo, 100x, 1.45 N.A. oil immersion, Nikon) was used for collecting fluorescence to maximize the photon collection efficiency. TIR was achieved by changing the angle of the incident laser on the back focal plane. The fluorescence emission from the sample passed through a dichroic (C-TIRF ultra high signal to noise 405/488/561/638 nm, Semrock) and band pass emission filters for Cy3 (Cat. No. ET607/36, Brightline, Semrock). Finally, fluorescence images were recorded on a scientific cMOS camera (Prime95B, Photometrix). For each sample, transfected neurons were located using low laser power (∼50 W/cm^2^) to minimize sample bleaching. 1000 frames were acquired for a 512 by 512-pixel region with an exposure time of 20 ms and without any delay between the frames. Analyses of TIR data were performed using Thunderstorm [61] and Trackmate [25] plugin in Fiji followed by plotting of diffusion coefficients using bespoke Python codes.

### 2.7 Proximity ligation assay

The proximity ligation assay (PLA) was performed as described previously [52; 55; 57] to visualize protein– protein interactions by microscopy. This assay is based on paired complementary oligonucleotide-labelled secondary antibodies that can hybridize and amplify a red fluorescent signal only when bound to two corresponding primary antibodies whose targets are in close proximity (within 40 nm). Briefly, trigeminal ganglion neurons were incubated overnight with 0.1% DMSO (as control) or **194** (5 µM). The next day, neurons were fixed using ice-cold methanol for 5 minutes and allowed to dry at room temperature.

The proximity ligation assay was performed according to the manufacturer’s protocol using the Duolink Detection Kit with PLA PLUS and MINUS probes for mouse and rabbit antibodies (Duolink in situ detection reagents red, cat. no. DUO92008; Duolink in situ PLA probe anti-rabbit MINUS, cat. no. DUO92005; Duolink in situ PLA probe anti-mouse PLUS, cat. no. DUO92001, Sigma-Aldrich). Primary antibodies (1/1000 dilution) were incubated for 1 hour at RT; Na_V_1.7 (cat. no. MABN41; Millipore, Research Resource Identifiers (RRID):AB_10808664), CRMP2 (cat. no. C2993; Sigma-Aldrich, RRID:AB_1078573), SUMO1 (cat. no. S8070; Sigma-Aldrich, RRID:AB_477543) and CRMP2 (cat. no. 11096; Tecan, immunobiological lab, RRID:AB_494511). Cells were then incubated overnight with anti-NeuN antibody (1/500; Cat #: D3S3I, Cell Signaling, RRID:AB_2630395) in PBS with 3% bovine serum albumin at 4°C. Cells were then washed 3 times in PBS and incubated with PBS containing 3% bovine serum albumin and secondary antibody, Alexa 488 goat anti-rabbit (Thermo Fisher Scientific, Waltham, MA) for 1 hour at room temperature. After washing with PBS, cells were stained with 49,6-diamidino-2-phenylindole (DAPI, 50 mg/mL) to detect cell nuclei and mounted in ProLong Diamond Antifade Mountant (cat. no. P36961, Life Technologies Corporation). Immunofluorescent micrographs were acquired using a Plan-Apochromat 63x/1.4 oil CS2 objective on a Leica SP8 confocal microscope operated by the LAS X microscope software (Leica). Camera gain and other relevant settings were kept constant throughout imaging sessions. Image J was used to count the number of PLA puncta per cell by an experimenter blinded to the treatment condition.

### 2.8 Patch-clamp electrophysiology

Whole-cell voltage-clamp and current-clamp recordings were performed between 18 h after culture or 36 h after transfection at room temperature by using an EPC 10 HEKA amplifier. In experiments where CRMP2 SUMOylation was prevented by the addition of 5 μM **194**, the compound was incubated in the tissue culture wells overnight, ∼14 h before the experiment. In experiments in which Na_V_1.7 channels were blocked with 5 nM ProTxII, the compound was added to the recording external solution.

#### 2.8.1 Voltage-clamp recordings

Trigeminal ganglion neurons were subjected to current-voltage (I-V) and activation/inactivation voltage protocols as follows: a) for the I-V protocol, cells were held at a potential of −60 mV and depolarized by 150-ms voltage steps from −70 mV to +60 mV in 5-mV increments, the resulting currents were normalized to the cell size, expressed as cell capacitance in pF, to get the corresponding current density and also infer the peak current density.

The activation voltage dependence of sodium channels was analyzed as a function of current versus voltage; b) inactivation protocol: from a holding potential of −60 mV, cells were subjected to 1-s hyperpolarizing/repolarizing pulses between−120 and 10 mV (+10 mV steps) followed by a 200-ms test pulse to +10 mV. This incremental increase in membrane potential conditioned various proportions of sodium channels into a state of fast inactivation. The external solution for voltage-clamp sodium recordings contained 140 mM NaCl, 30 mM tetraethylammonium chloride, 10 mM D-glucose, 3 mM KCl, 1 mM CaCl_2_, 0.5 mM CdCl_2_, 1 mM MgCl_2_, and 10 mM HEPES (pH 7.3 and 310-315 mOsm); and the internal solution consisted of: 140 mM CsF, 10 mM NaCl, 1.1 mM Cs-EGTA, and 15 mM HEPES (pH 7.3 and 290-310 mOsm). Analysis was performed by using Fitmaster software (HEKA).

For I−V curves, functions were fitted to data using a non-linear least-squares analysis. I−V curves were fitted using a double Boltzmann function:

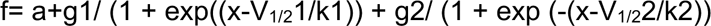

where x is the prepulse potential, V_1/2_ is the midpoint potential, and *k* is the corresponding slope factor for single Boltzmann functions. Double Boltzmann fits were used to describe the shape of the curves, not to imply the existence of separate channel populations. Numbers 1 and 2 simply indicate the first and second midpoints; *a* along with *g* are fitting parameters.

Activation curves were obtained from the I−V curves by dividing the peak current at each depolarizing step by the driving force according to the equation: G= I/ (V_mem_ − E_rev_), where I is the peak current, V_mem_ is the membrane potential, and Er_ev_ is the reversal potential.

The conductance (G) was normalized against the maximum conductance (G_max_). Inactivation curves were obtained by dividing the peak current recorded at the test pulse by the maximum current (I_max_). Activation and inactivation curves were fitted with the Boltzmann equation.

#### 2.8.2 Current-clamp recordings

Trigeminal ganglion neurons were held at their resting membrane potential and then subjected to a 100-millisecond depolarizing current step. The intensity of this current step was adjusted until the neurons fired an action potential to determine the rheobase of the cells. Additionally, depolarizing current injections of 0– 120 pA were applied to trigeminal ganglion cells to collect data on relative excitability, analyzed by counting the number of evoked action potentials during a 300-millisecond current step.

For whole-cell current-clamp experiments, the external solution contained 154 mM NaCl, 5.6 mM KCl, 2 mM CaCl_2_, 1 mM MgCl_2_, 10 mM D-glucose and 8 mM HEPES (pH 7.4 and 310–315 mOsm); and the internal solution consisted of: 137 mM KCl, 10 mM NaCl, 1 mM MgCl_2_, 1 mM EGTA, and 10 mM HEPES (pH 7.3 and 290–310 mOsm). The counting of the number of action potentials was performed by using Easy Electrophysiology software. A waveform analysis was carried out to calculate rise time, amplitude, and threshold by utilizing Easy Electrophysiology software.

Electrodes were pulled from filamented borosilicate glass capillaries (Warner Instruments) with a P-97 electrode puller (Sutter Instruments) to final resistances of 2.5–3.5 MΩ when filled with internal solutions. Whole-cell capacitance and series resistance were compensated.

Linear leak currents were digitally subtracted by the P/4 method for voltage clamp experiments, and bridge balance was compensated in current clamp experiments.

Signals were filtered at 10 kHz and digitized at 10–20 kHz. Cells in which series resistance or bridge balance was more than 15 MΩ or fluctuated by more than 30% over the course of an experiment were omitted from datasets.

### 2.9 Surgical Procedures

#### 2.9.1 Anesthesia

Rats of both sexes were anesthetized with intraperitoneal injections of ketamine (50mg/kg)/xylazine (7.5mg/kg) solution prior to any surgical procedure. A single investigator (OAK) performed the surgeries to minimize variability.

#### 2.9.2 ION-CCI surgery

Unilateral chronic constriction injury to the infraorbital nerve was performed to induce trigeminal neuropathic pain in rats as previously described [9; 13; 20; 22; 31; 36; 38; 40; 44; 49; 51-55; 57; 60; 63-65; 70; 71; 73; 74; 81; 82]. Briefly, following anesthesia, an approximately 1cm-long incision was made along the left gingivobuccal sulcus beginning just proximal to the first molar. The infraorbital nerve (ION) was exposed (∼0.5 cm) and freed from the surrounding tissue. Two chromic gut (4-0) ligatures were loosely tied around the exposed nerve. The incision was closed with absorbable sutures.

### 2.10 Intranasal compound administration

Twenty-five days post-ION-CCI, one group of rats received compound **194** intranasally (200 µg in 20 µL isotonic saline) whereas another group received 20 µL of isotonic saline (vehicle-control). Intranasal delivery was performed with a pipette and a disposable plastic tip. Immediately after administration, the head of the animal was held in a tilted back position for ∼15 seconds to prevent loss of solution from the nares. Behavioral assessments were done at 30 minutes, 1-, 2-, and 3-hours post-administration.

### 2.11 Assessment of somatosensory pain-like behavior

Von Frey detection threshold and pinprick response score assays were carried out to measure the hypersensitivity to mechanical stimulation applied over the ION dermatome. Somatosensory assessments were performed by a single investigator blinded to the experimental conditions.

Mechanical allodynia was measured by applying von Frey monofilaments delivering a calibrated amount of force ranging from 0.0008g to 1g or converted to log units 1.65 to 4.08 (EXACTA Precision & Performance monofilaments, Stoelting) within the ION dermatome in ascending order of intensity [45]. The lowest filament that evoked at least one withdrawal response was designated as the withdrawal threshold. A decrease in the withdrawal threshold is indicative of development of hypersensitivity.

Pinprick test was measured by scoring the response to stimulation with a blunted acupuncture needle applied within the vibrissal pad of rats. The scores were assigned as follows: 0 = no response, 1 = non-aversive response, 2 = mild aversive response, 3 = strong aversive response, 4 = prolonged aversive behavior [6; 76]. An increase in the response score is indicative of development of hypersensitivity.

### 2.12 Data analysis

All data sets corresponding to the electrophysiological, surface expression, PLA and *in situ* experiments were graphed and analyzed with GraphPad Prism (Version 9) [4]. They were checked for normality using the D’Agostino−Pearson test. Details of statistical tests, significance, and sample sizes are reported in the appropriate figure legends. All data plotted represent mean ± standard error of the mean (SEM). The statistical significance of differences between means was calculated by performing nonparametric Mann Whitney test or parametric analysis of variance (ANOVA) with a post hoc comparisons test (Tukey).

The statistical analyses for the behavioral data were performed using JMP Pro 16.0.0 software (JMP®, Version 16.0.0 SAS Institute Inc., 1989–2007). Goodness-of-fit test was used to evaluate normality of data. Analysis of variance was used to measure the between groups variability in normally distributed data and Wilcoxon/Kruskal Wallis Rank Sum’s nonparametric test was used to measure between groups differences of not normally distributed data. Results are reported as mean ± SEM unless otherwise specified. Wilcoxon Signed Rank test, a non-parametric alternative to paired *t*-test, was used to measure development of hypersensitivity following ION-CCI.

## 3. Results

### 3.1 Molecular identification and characterization of Na_V_1.7 neurons in the rat trigeminal ganglion

The expression and function of Na_V_1.7 channels as well as CRMP2 in rat TG neurons have been previously demonstrated [54; 81; 82]. Nevertheless, a comprenhensive search on their co-localization and the identity of the trigeminal nociceptor populations where they can be found has not been performed. We utilized *in situ* hybridization to characterize *Scn9a*-expressing TG neurons (defined by a DAPI-labeled nucleus surrounded by *Rbfox3* signal), using *Nefh* as a marker for A-∂ nociceptors and *Calca* to identify peptidergic C nociceptors (Figure 1A-B). Approximately 73% of all TG neurons were *Scn9a* positive, suggesting that 3/4 of TG neurons express the Na_V_1.7 channel (Figure 1C). Of *Scn9a* labeled neurons, approximately 68% were co-labeled for *Calca* and approximately 77% were co-labeled for *Nefh* (Figure 1D-E), indicating that *Scn9a* is highly expressed in both peptidergic C and A-∂ neurons.

**Figure 1.**
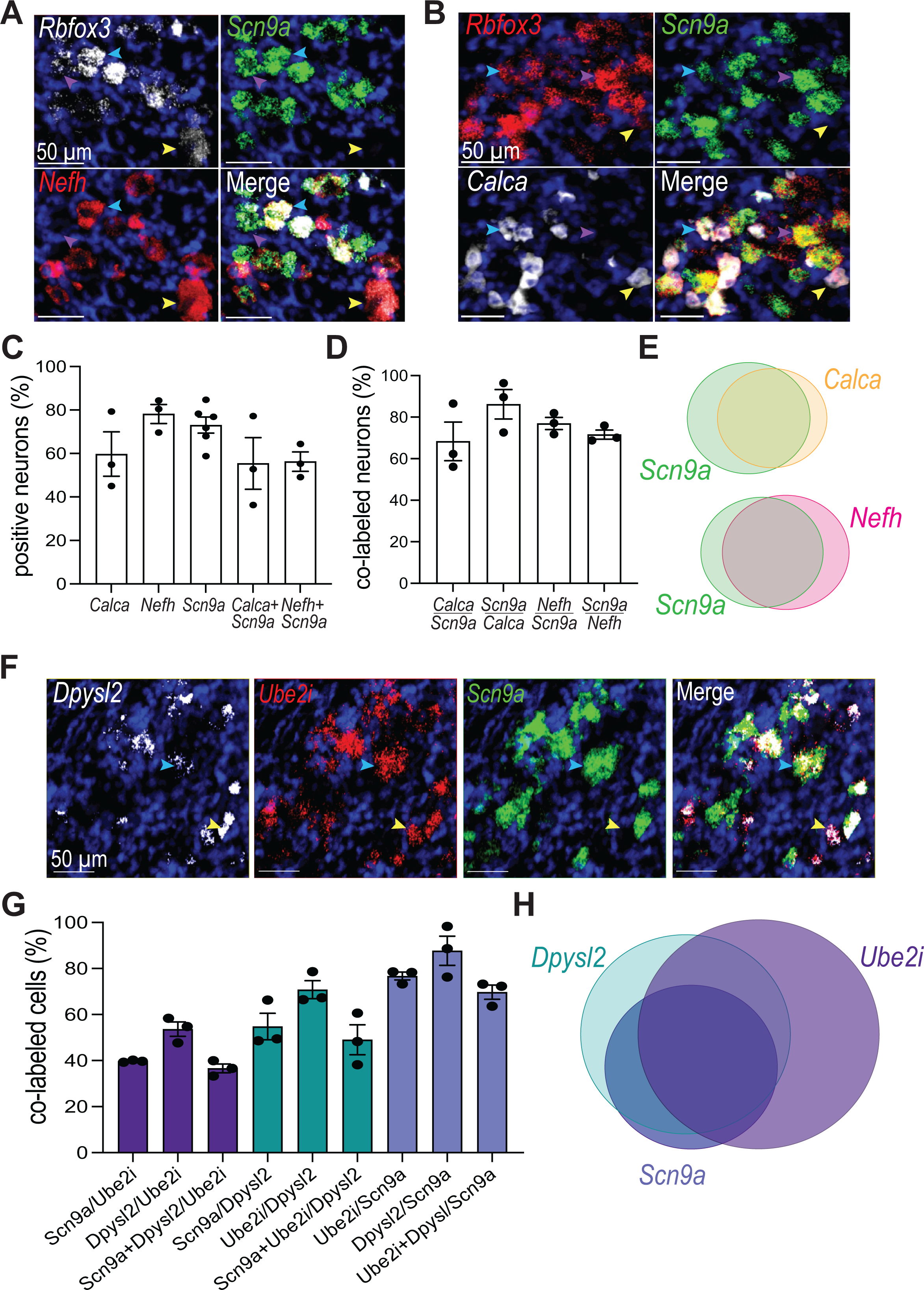
Characterization of *Scn9a* cells in rat trigeminal ganglion. Representative expression of *Scn9a* with *Nefh* (**A**) and Calca (**B**) in trigeminal ganglion neurons. Yellow arrowheads represent Scn9a negative *Nefh* or *Calca* neurons, purple arrowheads represent *Nefh* or *Calca* negative *Scn9a* neurons, blue arrowheads represent Scn9a neurons co-expressing either *Nefh* or *Calca*. **C.** Percentage of all neurons in the trigeminal ganglion expressing Calca, *Nefh*, *Scn9a*, or a combination of *Scn9a* and *Calca* or *Nefh*. **D.** Percent of *Scn9a*, *Calca*, or *Nefh* neurons co-expressing other markers. **E.** Visual representation of *Scn9a* co-expression with Calca or Nefh. **F.** Representative expression of *Dpysl2*, *Ube2i*, and *Scn9a* in the trigeminal ganglion. Yellow arrowheads represent *Scn9a* negative cells co-expressing Dpysl2 and Ube2i, blue arrowheads represent cells co-expressing *Dpysl2*, *Ube2i*, and *Scn9a*. **G.** Quantified percentages of *Ube2i*, *Dpysl2*, and *Scn9a* cells co-expressing other markers in the trigeminal ganglion. **H.** Visual representation of co-expression of *Dpysl2*, *Ube2i*, and *Scn9a*. All scale bars represent 50 µm. Each individual data point represents the mean of 4-6 sections from one animal. N= 3 to 6 rats per condition.

In order to explore the incidence of Na_V_1.7 with CRMP2 and the SUMO-conjugating enzyme Ubc9, we performed *in situ* hybridization in rat TG tissue for *Scn9a*, *Dpysl2*, and *UBE2I* (Figure 1F). *Scn9a* positive cells expressed high levels of both *Dpysl2* abd *UBE2I* (approximately 70%), suggesting that the majority of Na_V_1.7 TG neurons can be potentially regulated by CRMP2 SUMOylation (Figure 1G-H). Further, only 36% of *UBE2I* labeled cells expressed both *Scn9a* and *Dpysl2* and 50% and *Dpysl2* labeled cells expressed both *Scn9a* and *UBE2I* (Figure 1G-H), indicating that both *UBE2I* and *Dpysl2* are highly expressed in *Scn9a* negative neurons as well and also function in other TG populations.

### 3.2 CRMP2 and Na_V_1.7 physically interact in rat TG neurons and the small molecule 194 can uncouple this interaction

The SUMOylation of CRMP2 is a postranslational modification that modifies the surface expression, function, and promotes internalization of Na_V_1.7 channels [20; 22; 33; 52; 53]. Inhibition of this mechanism, both pharmacologically and genetically, has been demonstrated to promote the endocytosis of Na_V_1.7 in DRG neurons, reduce the excitability of dorsal horn neurons, and exhibit antinociceptive effects in animal pain models [20; 22; 33; 52; 53]. In the TG, however, no molecular correlation has been established between these two proteins. To address this, we employed proximity ligation assay (PLA) to examine the interactions between Na_V_1.7 and CRMP2, determine the SUMOylation status of CRMP2, and assess the disruptive effect of these two biological functions by compound **194** in TG neurons. PLA allows detection and quantification of protein complexes when proteins are within 40 nm of each other. The generation of a PLA signal can be interpreted as a protein-protein interaction. In cells treated with 0.1% DMSO, a positive PLA signal was found in TG neurons for SUMO1/CRMP2 and Na_V_1.7/CRMP2 (Figure 2A-D).

**Figure 2.**
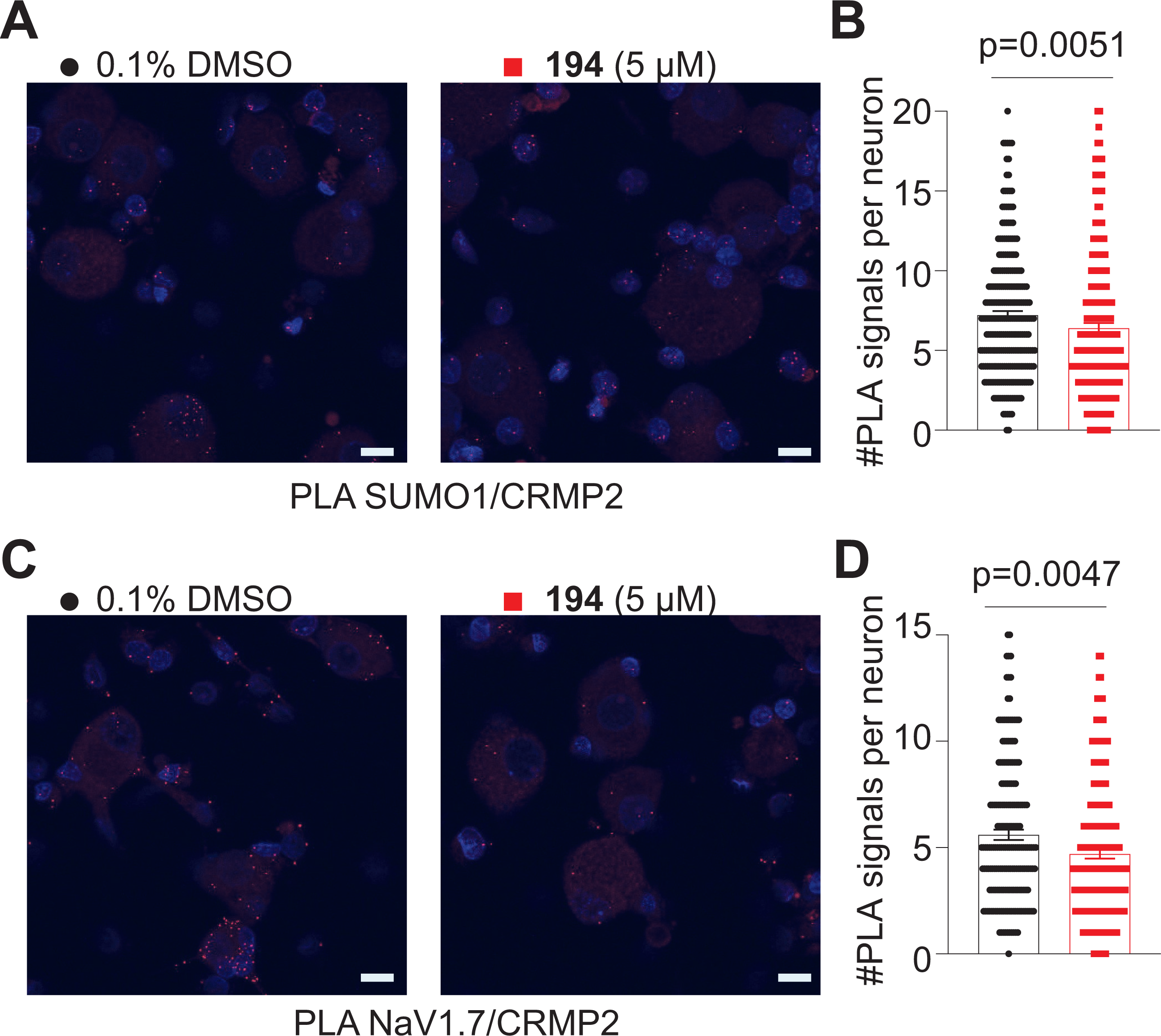
Compound 194 reduces CRMP2 SUMOylation decreasing binding of SUMO1 to CRMP2 and of Na_V_1.7 to CRMP2 in rat trigeminal ganglion (TG) neurons. Representative images of rat TG neuron cultures treated overnight with 0.1% DMSO (as a control) or 5 µM 194 following proximity ligation assay (PLA) between CRMP2 and SUMO1 (**A**), or between Na_V_1.7 and CRMP2 (**C**). NeuN signal was used to identify neurons. The PLA immunofluorescence labeled sites of interaction between CRMP2 and SUMO1 or Na_V_1.7 and CRMP2 (red puncta). In addition, nuclei are labeled with the nuclear labeling dye 4’,6’-diamidino-2-phenylindole (DAPI). Scale bar: 10 µm. Quantification of PLA puncta per neuron show that in TG neurons treated with **194**, SUMO1-CRMP2 (**B**) and Na_V_1.7-CRMP2 (**D**) interactions are significantly reduced compared with the control condition. p values as indicated; Mann Whitney test; Error bars indicate mean ± SEM.

Next, we tested if **194** could block the SUMOylation of CRMP2 and if this treatment could lead to Na_V_1.7/CRMP2 uncoupling. Overnight incubation of TGs with **194** significantly decreased both SUMO1/CRMP2 and Na_V_1.7/CRMP2 PLA signals compared to the control condition (0.1% DMSO; Figure 2A-D). These findings indicate that the interaction between Na_V_1.7 and CRMP2 is preserved in TG neurons and that **194** is able to disrupt this interaction by blocking the addition of SUMO1 to CRMP2 by SUMO conjugating enzyme Ubc9.

### 3.3 CRMP2 SUMOylation inhibitor compound 194 decreases the membrane expression, but increases clustering, of Na_V_1.7 channels in rat trigeminal ganglion (TG) neurons

Our group has reported that the CRMP2 SUMOylation inhibitor, compound **194**, can reduce the cell surface expression of Na_V_1.7 in DRG neurons, facilitating the internalization of the channel [13]. We used immunofluorescent labeling followed by confocal imaging to evaluate if the membrane expression of Na_V_1.7 is affected by **194 **treatment of cultured rat TG neurons. Results showed that the surface expression of this channel was decreased in cells treated with **194**, compared to the control condition (Figure 3A, B). These data suggest that **194** is promoting the internalization of Na_V_1.7 in rat TG neurons.

**Figure 3.**
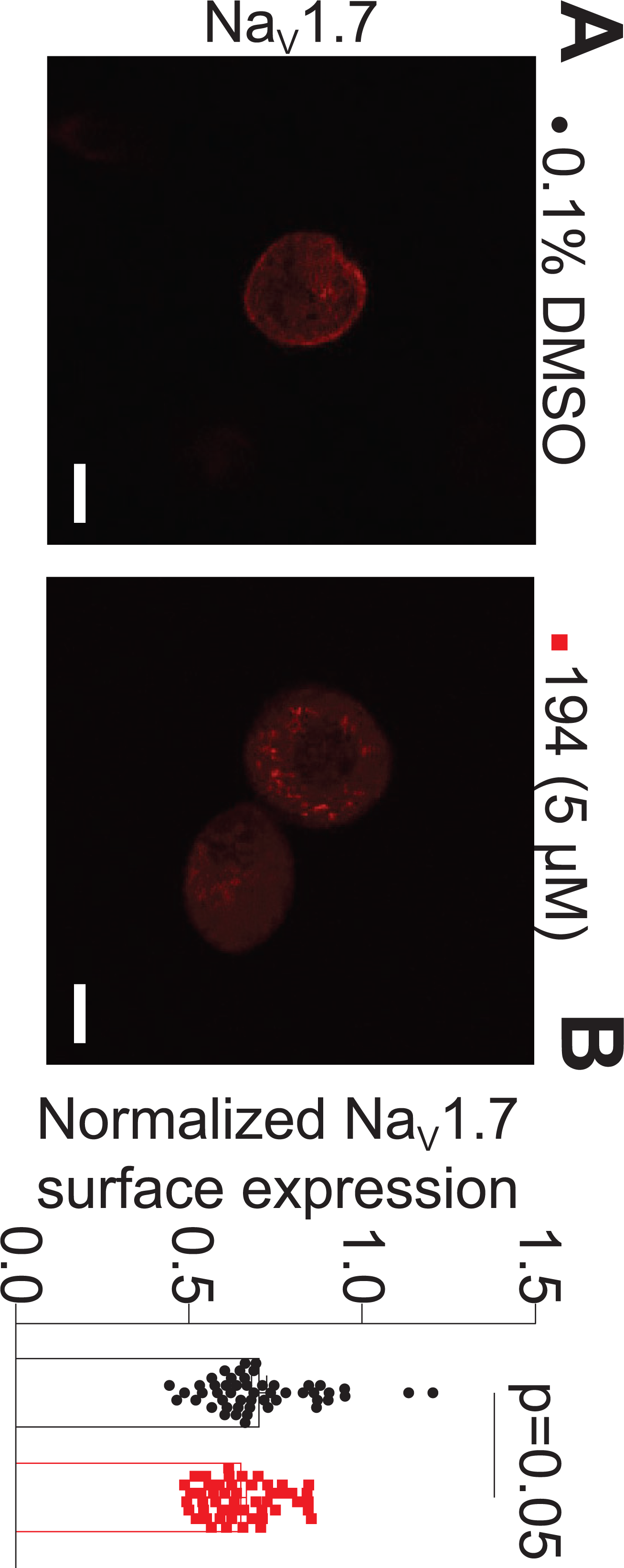
Compound 194 reduces CRMP2 SUMOylation causing a decrease in Na_V_1.7 surface expression in rat trigeminal ganglion (TG) neurons. **A** Representative confocal images of rat TG neuron cultures treated overnight with 0.1% DMSO (as a control) or 5 µM of 194 and labeled with an antibody against Na_V_1.7. Scale bar: 10 µm. **B** Quantification of normalized surface expression of NaV1.7 per neuron shows that TG neurons treated with 194 have decrease surface expression of Na_V_1.7 compared with control condition. n = 50-51 cells; p values as indicated; Unpaired t-test; error bars indicate mean ± SEM.

To independently corroborate this effect on surface expression of Na_V_1.7, we used super-resolution microscopy to resolve Na_V_1.7 channels at single molecule resolution. We fused a 29 kilodalton Photostability Enhancing Peptide (PEP) for the membrane impermeable dye cyanine (PEPCy) to the N-terminus of Na_V_1.7 (Figure 4A). To test the functionality of this channel, we trasfected HEK293 cells with Na_V_1.7-PEPCy and measured macroscopic sodium currents. Depolarizing pulses ranging from −70 to +60 mV produced a rapidly activating and inactivating family of sodium currents (Figure 4B) with a peak between −20 and 0 mV (Figure 4C) and a peak current density of −46.48 ± 9.74 pA/pF (n = 14; Figure 4D). Normalized conductance values were fit with the Boltzmann equation to determine the voltage dependence of activation. Half-activation value was −19.64 ± 1.34 (n=12) and (slope) *k* = 7.81 ± 1.33 (n=12)(Figure 4E). Steady-state fast-inactivation curves were generated by applying 500 ms inactivating potentials from −120 to −10 mV, followed by a 40 ms step to −10 mV. Normalized conductance values were fit with the Boltzmann equation to determine the voltage dependence of steady-state fast-inactivation. Half-inactivation value was: −75.18 ± 1.45 (n=12) and *k* = 9.85 ± 1.36 (n=12)(Figure 4E). The combined information affirms that the introduction of PEPCy3 into the channel did not affect its capability to generate functional Na_V_1.7 channels.

**Figure 4.**
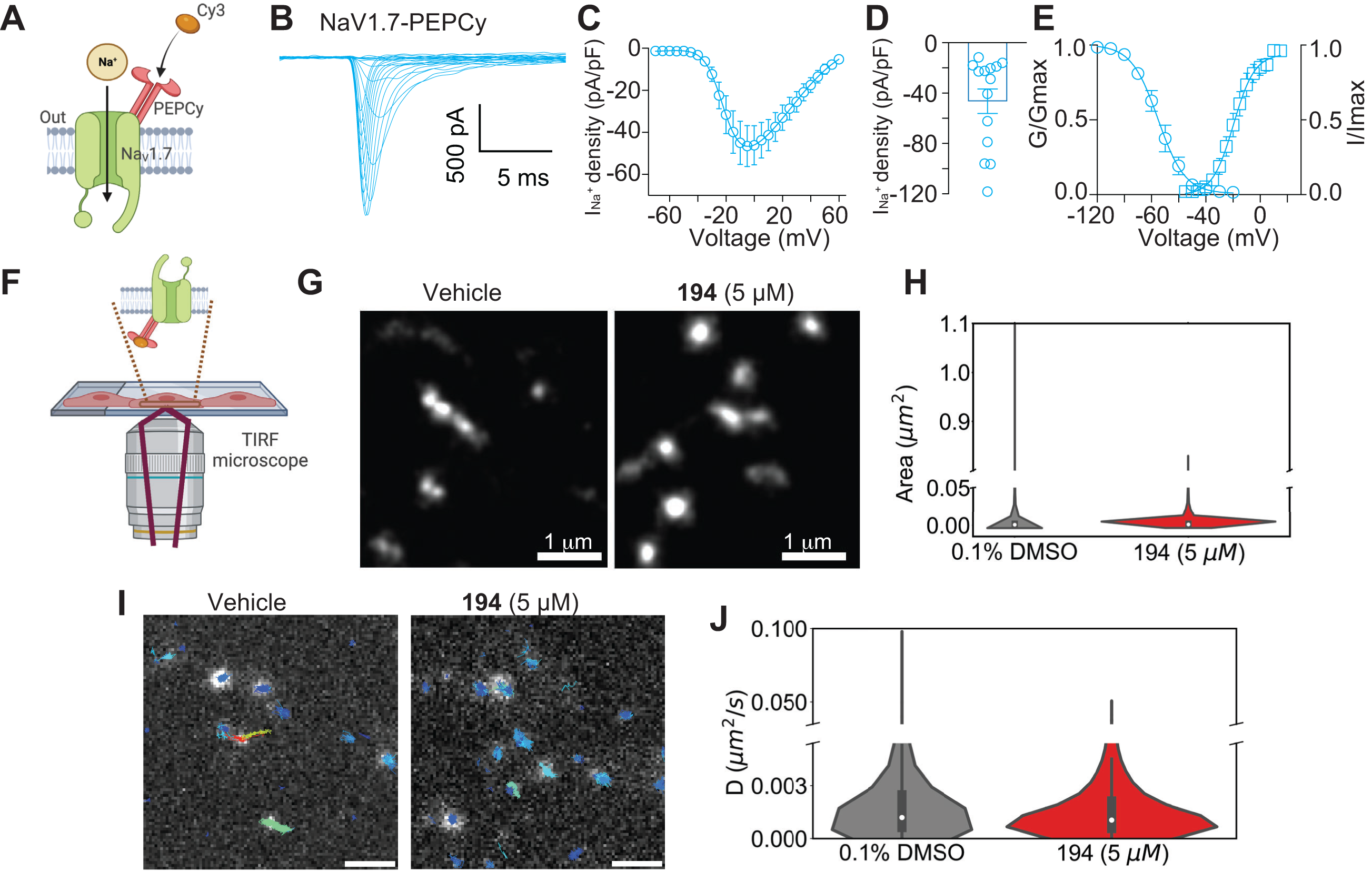
TIR microscopy reveals effects on reduced diffusivity of single Na_V_1.7 molecules upon treatment with compound 194. **A.** Schematic of the NaV1.7-PEPCy construct and site of binding of cell impermeant Cy3 dye. The schematic was generated with BioRender. **B.** Representative traces of sodium currents (I_Na_^+^) recorded from HEK 293 cells transiently expressing Na 1.7-PEPCy and green fluorescent protein (GFP). Cells were identified with GFP fluorescence. **C.** Summary of I_Na_^+^ density versus voltage relationship. **D.** Bar graphs of total peak I_Na_^+^ density from HEK 293 cells transiently expressing Na 1.7-PEPCy. **E.** Boltzmann fits for normalized conductance voltage relationship for voltage dependent activation (*G/Gmax*) and inactivation (*I/Imax*) from HEK293 cells transiently expressing Na_V_1.7-PEPCy. Error bars indicate mean ± SEM; *n*= 12 cells. **F.** Schematic of Total Internal Reflection (TIR) Microscopy used for imaging TG neurons transfected with Na_V_1.7-PEPCy. **G.** Super-resolved images showing single molecule localizations in a representative field of TG neurons incubated without (*left*) or with (*right*) 5 µM of compound **194**. Without drug treatment single molecules of Na_V_1.7 assemble in large clusters and as mobile single molecules compared to more clustered Na_V_1.7 molecules in the presence of **194**. Scale bar is 1 µm. **H.** Distribution of cluster areas shown as a violin plot for control and **194** treated TG neuron samples. The white dot within the bars in the violin plot denotes the median of the distribution. Between 36000-50000 clusters were analyzed from fifteen regions for each distribution. **I.** Super-resolved images showing single molecules in first frame overlaid with their trajectories in a representative field of TG neurons incubated without (*left*) or with (*right*) 5 µM of compound **194**. Each trajectory is color-coded by track displacement in µm. Without drug treatment, single molecules of Na_V_1.7 are more diffusive compared to Na_V_1.7 molecules in the presence of **194**. Scale bar is 2 µm. **J.** Distribution of two-dimensional diffusion coefficients shown as a violin plot for control and **194** treated TG neuron samples. The white dot within the bars in the violin plot denotes the median of the distribution. Between 4200-4300 single molecules were analyzed from twelve regions for each distribution.

Total internal reflection (TIR) microscopy on TG neurons transfected with Na_V_1.7-PEPCy3 labeled with the membrane impermeable dye Cy3 revealed single molecules and clusters of Na_V_1.7 on the cell surface (Figure 4F). Single molecule localization analyses from super-resolution microscopy revealed the presence of clusters of various sizes (Figure 4G). Interestingly, the observed Na_V_1.7-PEPCy3 cluster sizes were dependent on the presence or absence of compound **194**. In the presence of **194** we observed a median cluster size of 0.0008 µm^2^ versus 0.003 µm^2^ in the untreated case (Figure 4H). A smaller cluster could arise from slow diffusion of molecules. To test this hypothesis, we performed a trajectory analysis of single NaV1.7-PEPCy3 proteins. Visual inspection of trajectories revealed that TG neurons transfected with Na_V_1.7-PEPCy3 and treated with **194** exhibited lower mobility compared to untreated cells (Figure 4I, Supplementary Videos 1 and 2). We analyzed the motion of single molecules from both samples using mean squared displacement (MSD) analysis [50]. Using the first three points of the MSD versus lag time plot, we computed the distribution of two-dimensional diffusion coefficients (D) for the trajectories. For the **194**-treated samples, the distribution of D had a median value of 0.0009 µm^2^/s, which was approximately two-fold lower than the median D for the untreated samples (D = 0.001 µm^2^/s) (Figure 4J). Additionally, the untreated samples exhibited many highly diffusive molecules (D values greater than 0.05 µm^2^/s)(Supplementary Video 1). Together, these data demonstrate that **194** treatment inhibits Na_V_1.7 diffusion on the membrane leading to the formation of tightly localized membrane clusters.

### 3.4 Preventing CRMP2 SUMOylation with compound 194 decreases the activity of Na_V_1.7 channels in rat trigeminal ganglion neurons

To assess the functional implications of CRMP2 SUMOylation on Na_V_1.7 channels in TG neurons, we initially examined the impact of **194** on total sodium currents. Sodium currents from TG cells held at −60 mV and depolarized by 150-ms voltage steps from −70 mV to +60 mV in 5-mV increments were normalized to the cell’s capacitance to obtain the current-voltage relationships shown in Figure 5. Large transient inward currents were seen in response to depolarizing steps (Figure 5A). Mean peak current density (at - 5 mV) of DMSO treated TG neurons was −525.3 ± 40.3 pA/pF (DMSO, n = 29, Figure 5B-C). Blocking CRMP2 deSUMOylation by overnight incubation with **194** (5 µM) decreased total sodium currents and current density, causing a significant diminution in peak sodium currents of ∼ 58% (−220.8 ± 29.8 pA/pF, n = 30, Figure 5A-C) with respect to control (i.e., DMSO-treated TG neurons). Analysis of voltage-dependence of activation revealed that treating TG neurons with **194** caused a significant ∼6 mV shift in the activation curve towards more positive potentials. Moreover, **194** induced a significant ∼5 mV leftward shift in the voltage-dependence of inactivation (Figure 5D and Table 1).

**Figure 5.**
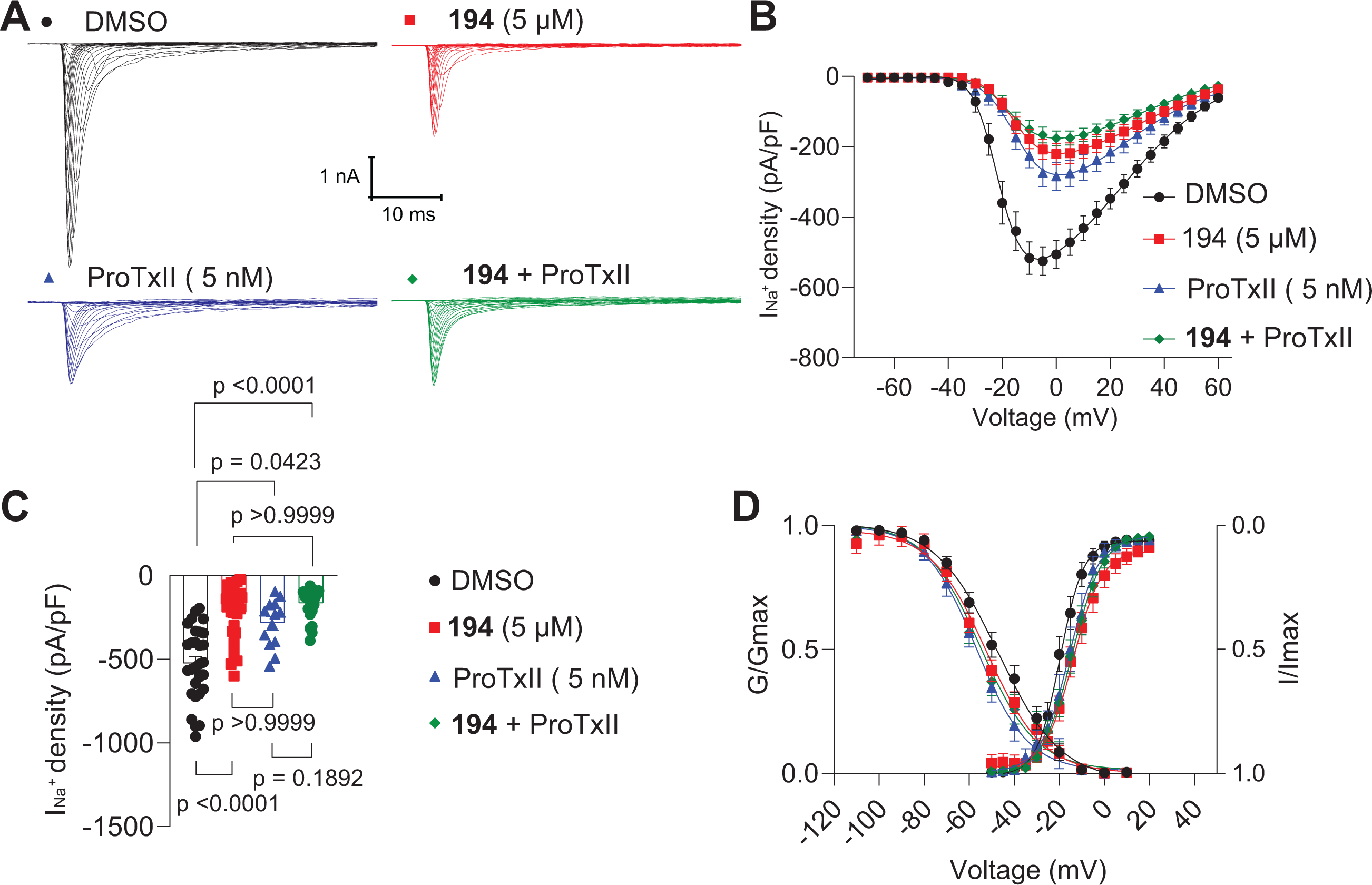
Blocking CRMP2 SUMOylation with compound 194 reduces sodium currents of rat trigeminal ganglion sensory neurons. **A.** Representative traces of sodium currents (I_Na_^+^) from rat trigeminal ganglion neurons (TGs) incubated with 0.1% DMSO (control; black circles), 5 µM **194** (red squares), 5 nM ProTxII (blue triangles) and 5 µM **194** + 5 nM ProTxII (green diamonds). **B.** Summary of total I_Na_^+^ density versus voltage relationship. **C.** Bar graphs of total peak I_Na_^+^ density from TGs pre-treated as indicated. Significant differences were observed in the I_Na_^+^ density of each group, when compared to the control (one-way ANOVA followed by a Tukey’s post hoc test, *n*= 14 to 30 cells per condition from four different animals). **D.** Boltzmann fits for normalized conductance voltage relationship for voltage dependent activation (*G/Gmax*) and inactivation (*I/Imax*) of rat TG neurons pre-treated as indicated. Significant differences were observed in the V_1/2act_ and V_1/2inact_ of each group, when compared to the control (one-way ANOVA followed by a Tukey’s post hoc test, *n*= 14 to 30 cells per condition). Error bars indicate mean ± SEM.

**Table 1.** Gating properties of voltage-gated sodium channels in rat trigeminal ganglion neurons.

| Condition | Activation |  | Inactivation |  |
| --- | --- | --- | --- | --- |
| | $V_{0.5}$ | $k$ | $V_{0.5}$ | $k$ |
| *Figure 5D* |  |  |  |  |
| DMSO | -19.626 ± 0.598<br>(29) | 5.201 ± 0.535<br>(29) | -47.260 ± 1.348<br>(29) | -13.943 ± 1.337<br>(29) |
| 194 (5 µM) | -13.742 ± 0.907<br>(30)<br>p < 0.0001 vs.<br>DMSO | 6.343 ± 0.841<br>(30) | -52.294 ± 1.463<br>(30)<br>p = 0.0440 vs.<br>DMSO | -12.521 ± 1.403<br>(30) |
| DMSO+<br>ProTx-II (5 nM) | -15.439 ± 1.021<br>(13)<br>p = 0.0040 vs.<br>DMSO | 6.446 ± 0.951<br>(13) | -56.828 ± 1.625<br>(13)<br>p = 0.0009 vs.<br>DMSO | -11.461 ± 1.513<br>(13) |
| **194** (5 µM) +<br>ProTx-II (5 nM) | -14.307 ± 0.722<br>(23)<br>p < 0.0001 vs.<br>DMSO | 6.854 ± 0.683<br>(23) | -54.979 ± 1.424<br>(23)<br>p = 0.0014 vs.<br>DMSO | -12.416 ± 1.357<br>(23) |
| *Figure 6D* |  |  |  |  |
| Scramble siRNA | -21.225 ± 0.565<br>(24) | 3.908 ± 0.494<br>(24) | -41.770 ± 1.722<br>(24) | -16.145 ± 1.758<br>(24) |
| CRMP2 siRNA | -21.519 ± 1.132<br>(35) | 7.718 ± 1.071<br>(35)<br>p = 0.0192 vs.<br>Scramble siRNA | -49.580 ± 1.648<br>(35)<br>p = 0.0072 vs.<br>Scramble siRNA | -17.560 ± 1.852<br>(35)<br>p=0.9369 |
| Scramble siRNA +<br>**194** (5 µM) | -17.052 ± 0.730<br>(18)<br>p = 0.0480 vs.<br>scramble siRNA | 5.941 ± 0.668<br>(18) | -54.172 ± 1.171<br>(18)<br>p = 0.0001 vs.<br>Scramble siRNA | -12.332 ± 1.115<br>(18) |
| CRMP2 siRNA +<br>**194** (5 µM) | -14.844 ± 1.261<br>(18)<br>p = 0.0007 vs.<br>Scramble siRNA | 8.155 ± 1.259<br>(18)<br>p = 0.0295 vs.<br>Scramble siRNA | -46.227 ± 2.414<br>(18) | -17.975 ± 2.662<br>(18) |

Values are means ± S.E.M. calculated from fits of the data from the indicated number (in parentheses) of cells to the Boltzmann equation. $V_{0.5}$ , midpoint potential (mV) for voltage-dependent activation or inactivation; $k$ , slope factor. Data were analyzed with a one-way ANOVA followed by a Tukey's post hoc test.

We next utilized Protoxin-II (ProTx-II), a selective Na_V_1.7 channel blocker [67], to assess the contribution of Na_V_1.7 channels to the reduction in total sodium currents caused by **194**. Acute application of ProTx-II revealed a similar decrease in sodium currents and current density (∼46%, −284 ± 39.5 pA/pF, n = 13, Figure 5A-C) to that observed in the presence of compound **194**. When analyzing the kinetics of activation and inactivation, we found that ProTxII promoted a significant ∼4 mV shift in the activation curve towards more positive potentials as well as a ∼9 mV leftward shift in the voltage-dependence of inactivation (Figure 5D and Table 1).

Next, to test if compound **194** and ProTx-II converge on modulating Na_V_1.7 channels, we performed oclussion experiments where we first incubated TG neurons with **194** and then applied ProTx-II. We found that co-application of both compounds did not cause a further reduction in sodium currents (∼ 66% respect to control, −175.6 ± 20.8 pA/pF, n = 23, Figure 5A-C) when compared to cells treated with **194** and ProTx-II alone. This suggests that Na_V_1.7 channels had already been maximally blocked/regulated. Importantly, no significant changes were observed in tetrodotoxin-resistant sodium channels —Na_V_1.8 and Na_V_1.9— in the presence of **194** (data not shown).

Overall, these data support the conclusion that modulation of Na_V_1.7 channels by CRMP2 SUMOylation occurs in TG neurons, and that sodium current reductions imposed by **194** are mainly due to reduced current influx through Na_V_1.7 channels.

### 3.5 CRMP2 knockdown decreases sodium currents and recapitulates the effect of 194 on rat trigeminal ganglion neurons

Our data thus far demonstrates that Na_V_1.7 channels represent an important component in rat TG neurons and that the SUMOylation of a cytosolic regulator CRMP2, can modify the function of these channels, ultimately regulating neuronal excitability. These observations position CRMP2 as a central driver of Na_V_1.7 activity and underscore the inhibitory effect exerted by compound **194** [20]. Considering that CRMP2’s loss of SUMOylation inhibits Na_V_1.7 activity, a similar reduction in Na_V_1.7 currents would be expected when CRMP2 expression is reduced. To test if CRMP2 is the target protein of **194**, we proceeded to evaluated if silencing CRMP2 in TG neurons would modify the effect of compound **194** on sodium current density. Firstly, we knocked-down the expression of CRMP2 in TG neurons by using a previously validated siRNA against CRMP2 [11; 20] and measured sodium currents by performing whole-cell voltage clamp electrophysiology. Silencing CRMP2 led to a significant reduction (∼35%) in peak sodium currents and current densities (−209.4 ± 21.5 pA/pF, n = 35, Figure 6A-C) when compared to the control condition (−323.2 ± 30.6 pA/pF, n = 24, Figure 6A-C). Analysis of voltage-dependence showed a significant ∼8-mV shift in the V_0.5_ of inactivation towards more negative potentials (Figure 6D). Overnight incubation of **194** alone (5 µM) caused a significant reduction of sodium currents (∼ 53%, −150.5 ± 24.4 pA/pF, n = 18,) which did not decrease further when the compound was applied to CRMP2-silenced cells (∼ 57%, −136.9 ± 21.6 pA/pF, n = 18) (Figure 6A-C). Compound **194** promoted a rightward shift (∼4 mV) in the V_0.5_ of activation of control cells and a ∼13-mV shift in the V_0.5_ of inactivation towards more negative voltages (Figure 6D, Table 1). A ∼7-mV rightward shift in the V_0.5_ of activation was observed when **194** was added to the CRMP2-silenced cells (Figure 6D, Table 1). These data functionally validate the role of CRMP2 in modulating Na_V_1.7 function in TG neurons. Furthermore, our findings suggest that the target effector protein of compound **194** is indeed CRMP2, which subsequently allosterically modulates Na_V_1.7 channels.

**Figure 6.**
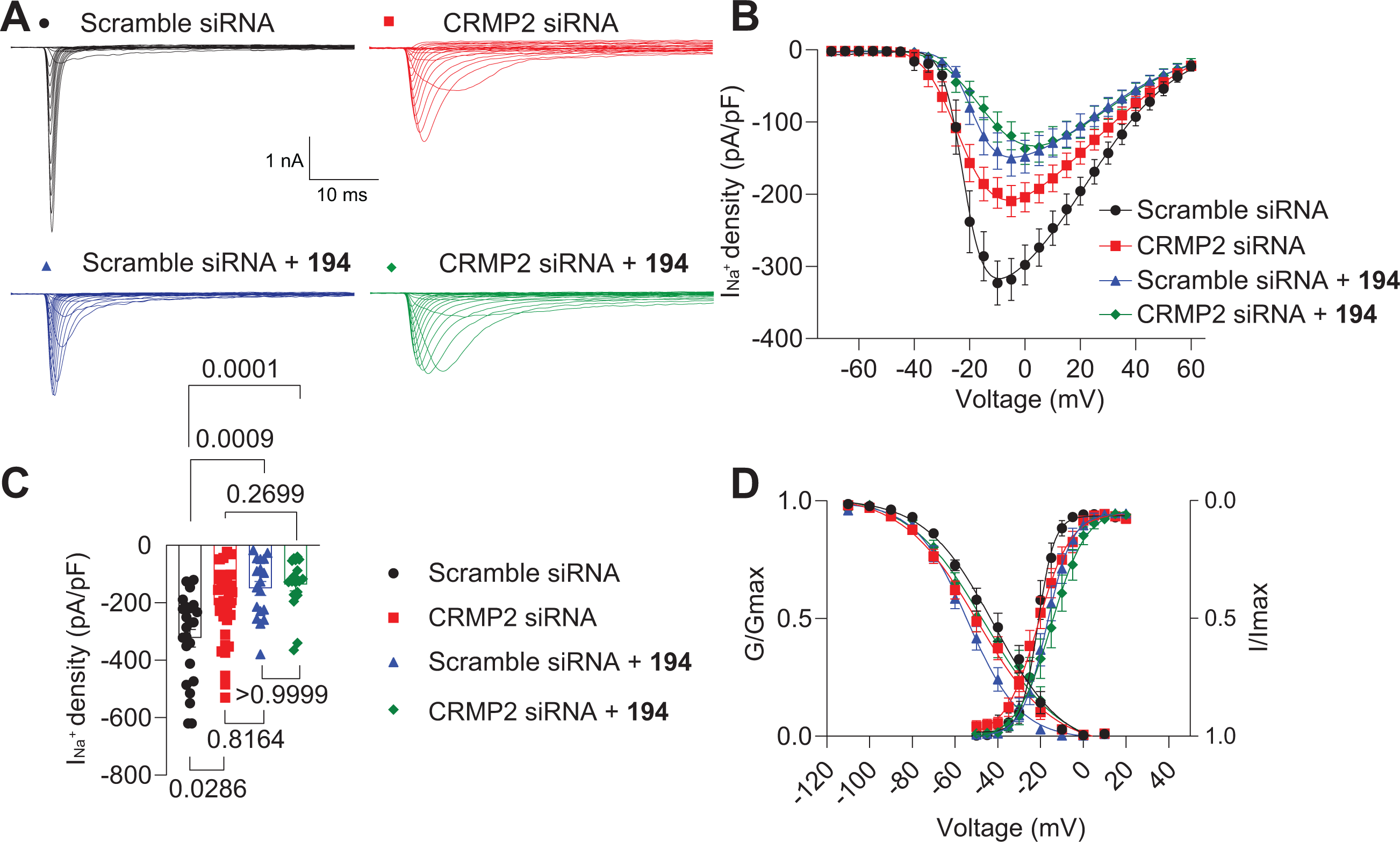
Knockdown of CRMP2 reduces sodium currents in rat trigeminal ganglion (TG) sensory neurons to a similar extent as compound 194. **A.** Representative traces of sodium currents (I_Na_^+^) from rat TG neurons transfected with scramble siRNA (black circles), CRMP2 siRNA (red squares), scramble siRNA + **194** (blue triangles), or CRMP2 siRNA + 194 (green diamonds). **B.** Summary of total I_Na_^+^ density versus voltage relationship. **C.** Bar graphs of total peak I_Na_^+^ density from TGs pre-treated as indicated. Significant differences were observed in the V_1/2act_ of each group, when compared to the control (one-way ANOVA followed by a Tukey’s post hoc test, *n*= 18 to 35 cells per condition). **D.** Boltzmann fits for normalized conductance voltage relationship for voltage dependent activation (*G/Gmax*) and inactivation (*I/Imax*) of rat TG neurons pre-treated as indicated. Significant differences were observed in the V_1/2act_ and V_1/2inact_ of each group (Table 1), when compared to the control (one-way ANOVA followed by a Tukey’s post hoc test, *n*= 18 to 35 cells per condition from four different animals). Error bars indicate mean ± SEM.

### 3.6 The CRMP2 SUMOylation inhibitor 194 decreases excitability of rat trigeminal ganglion neurons

As Na_V_1.7 channels play a crucial role in determining the threshold for action potential firing in sensory neurons [48; 77], the observed regulation of their activity in our voltage-clamp experiments is likely to have a direct impact on neuronal excitability. To test if preventing the SUMOylation of CRMP2 and resulting Na_V_1.7 inhibition have an impact on the excitability properties of TG neurons, we performed current-clamp experiments in the absence and presence of **194**. We observed that overnight incubation of TG neurons with 5 µM of **194** significantly decreased the number of action potentials from 80 to120 pA-current injection steps (Figure 7A, B). When analyzing the maximum number of action potentials, quantified at 120 pA, we observed a ∼50% decrease in the presence of **194** (3.23 ± 2.029) compared to the control group (6.4 ± 3.544; Figure 7A, B). No changes were observed in the rheobase, which is the minimum current required to evoke a single action potential, or in the basal resting membrane potential (Figure 7C, D). A waveform analysis of the evoked action potentials revealed that **194** significantly increases the rise time to peak and the threshold at every current step while, in turn, the amplitude was found to be decreased (Figure 7E-G). Collectively, these findings strongly indicate that **194** exerts a significant influence on the excitability of TG neurons, primarily by inhibiting Na_V_1.7 activity through the reduction of CRMP2 SUMOylation.

**Figure 7.**
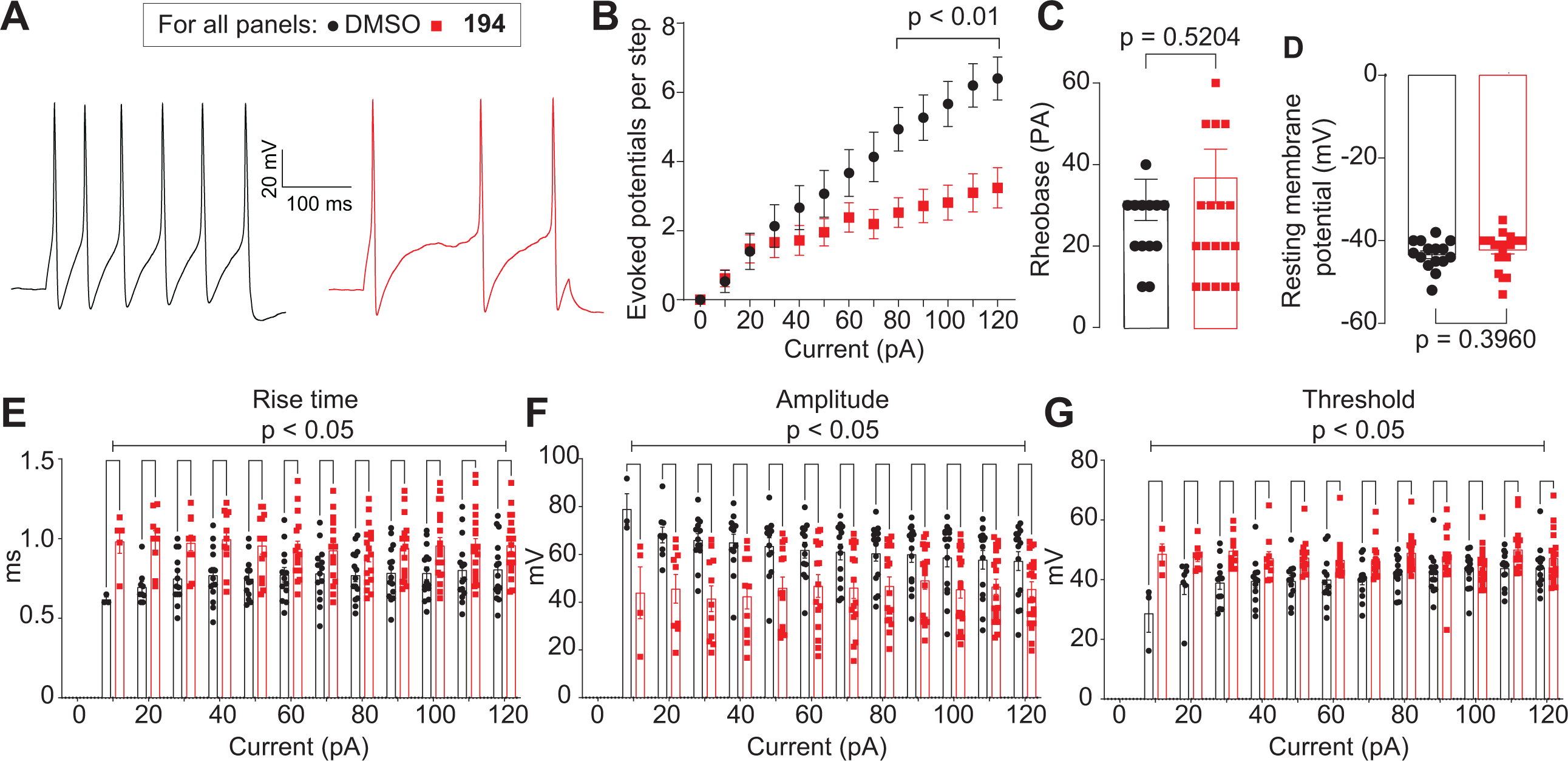
Preventing CRMP2 SUMOylation with compound 194 decreases excitability of rat trigeminal ganglion sensory neurons. **A** Representative traces of evoked action potentials at 120 pA from rat trigeminal ganglion neurons (TGs) incubated overnight with 0.1% DMSO (control; black circles) or 5 µM **194** (red squares). **B** Summary of the current-evoked action potentials in response to current injections between 0-120 pA in the presence of overnight 0.1% DMSO or 5 µM **194**. **C** Quantification of the rheobase in the presence of overnight 0.1% DMSO or 5 µM **194 D** Quantification of the resting membrane potential in the presence of overnight 0.1% DMSO or 5 µM **194**. Waveform parameters, (**E**) rise time (in ms), (**F**) amplitude (in mV) and (**G**) threshold (in mV) of such evoked action potentials. Unpaired single and multiple *t* tests, *n*= 15 to 21 cells per condition from four different animals. Error bars indicate mean ± SEM.

### 3.7 Intranasal administration of CRMP2 SUMOylation inhibitor 194 decreases unilateral ION-CCI-mediated mechanical allodynia in rats

Our previous study demonstrated that the small molecule **194** effectively reverses pain in six different models across two rodent species, administered via four routes [13]. However, treating craniofacial pain, like PTNP, poses challenges as common drug delivery methods may not effectively reach cephalic areas to alleviate pain [8; 17]. To manage trigeminal pain, intranasal administration offers a non-invasive drug delivery route, where drugs are absorbed by the nasal cavity’s mucus membrane, reaching systemic blood circulation and trigeminal nerves [8; 17]. Given **194**’s proven *in vitro* effects on rat trigeminal Na_V_1.7 channels and excitability, we investigated its potential for ameliorating trigeminal neuropathic pain through intranasal delivery in rats.

#### 3.7.1 Establishment of the unilateral ION-CCI-mediated mechanical allodynia model in rats

We first measured the development of mechanical allodynia at 17-, 21- and 24-days post ION-CCI. Male and female animals developed significant hypersensitivity as indicated by a significant drop in von Frey detection thresholds (vF) at 17 (1.65 ± 0.0 for males and females), 21 (1.65 ± 0.0, for male and females), and 24 (1.65 ± 0.0, for male and females) days following ION-CCI compared to baseline (2.68 ± 0.28 for males and 2.43 ± 0.01 for females) (Figures 8A, B). Furthermore, there was a significant increase in pin prick (PP) score at 17 (2.90 ± 0.45 for males and 3.00 ± 0.00 for females), 21 (2.85 ± 0.49 for males and 3.00 ± 0.00 for females), and 24 (2.8 ± 0.41 for males and 3.00 ± 0.00 for females) days following ION-CCI compared to baseline (1.25 ± 0.44 for males and 1.60 ± 0.18 for females) (Figures 8C, D). These behavioral results show that both male and female animals exhibited significant hypersensitivity following ION-CCI.

**Figure 8.**
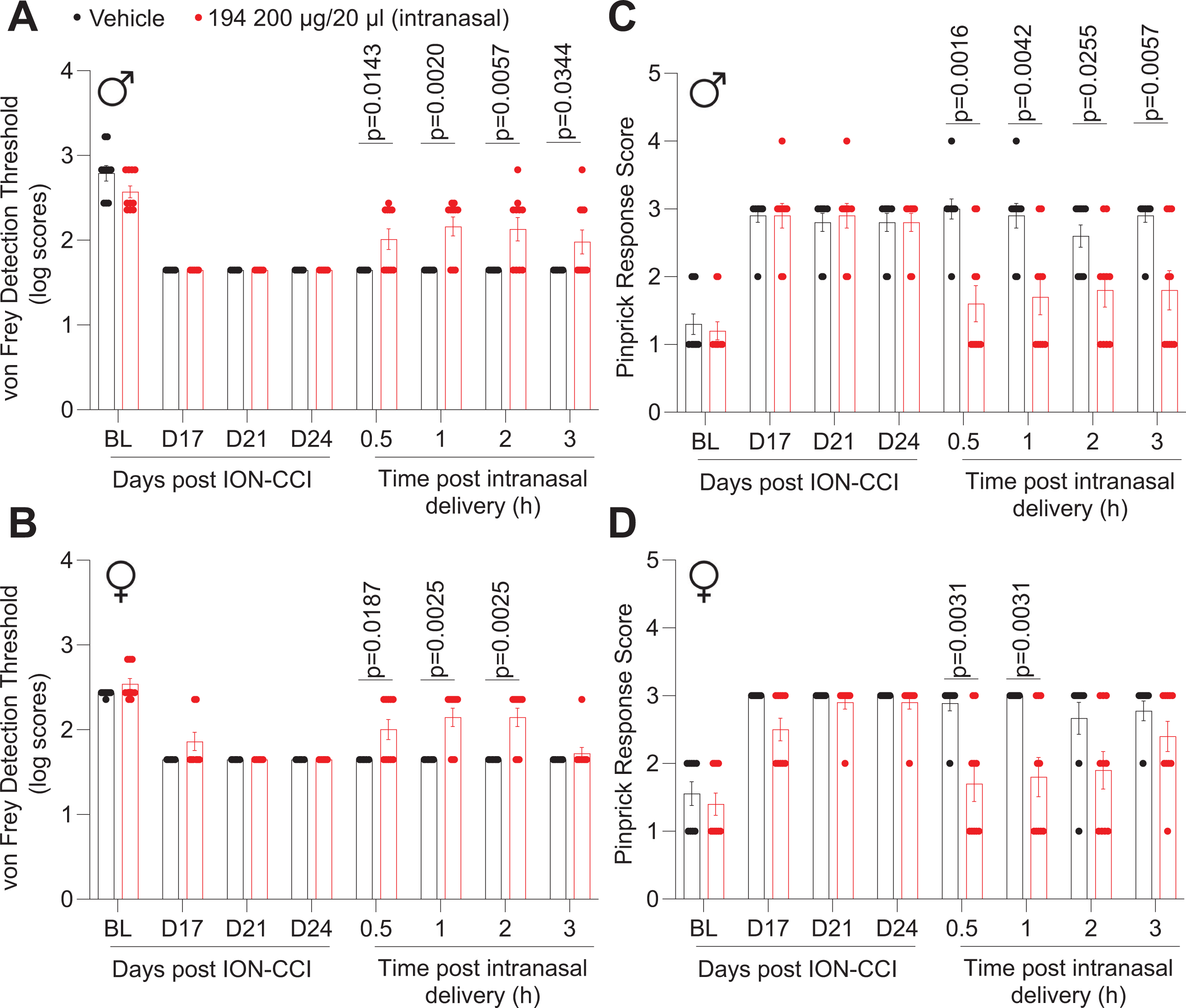
Intranasally-administered CRMP2 SUMOylation inhibitor 194 decreases mechanical hypersensitivity in rats with trigeminal neuropathic pain. **A.** Timeline of the experimental paradigm indicating that rats developed significant hypersensitivity to von Frey monofilaments at days 17, 21 and 24 post chronic constriction of the infraorbital nerve (ION-CCI) compared to baseline (BL). Intranasal compound 194 resulted in a significant increase in the von Frey detection threshold in the ipsilateral side of ION-CCI rats compared to ION-CCI rats who received vehicle (saline). Von Frey detection threshold was evaluated at 30 min,1-, 2- and 3-hours post compound 194 administration. p value as indicated; Steel-Dwass test. N = 9 −10 rats per treatment; error bars indicate mean ± SEM. **B.** Timeline of the experimental paradigm indicating that rats developed significant hypersensitivity as indicated by the increase in pinprick response scores at days 17, 21 and 24 post ION-CCI compared to baseline (BL). Pinprick response scores were observed at 30 min,1-, 2- and 3-hours post compound 194 administration. p value as indicated; Steel-Dwass test. N= 9 −10 rats per treatment. Error bars indicate mean ± SEM.

#### 3.7.2 Effects of intranasal administration of compound **194** on von Frey detection thresholds in rats with ION-CCI

Intranasally delivery of compound **194** (200 µg / 20 µL) effectively increased mechanical detection threshold in the side of the face ipsilateral to ION-CCI compared to vehicle (i.e., saline) treated animals. A significant increase in detection threshold was observed beginning at 30 min post administration (194: 2.01 ± 0.38, saline 1.65 ± 0.00), and the effect remained significant at 1 hour (194: 2.16 ± 0.36, saline 1.65 ± 0.00), 2 hours (194: 2.13 ± 0.44, saline 1.65 ± 0.00) and 3 hours (194: 1.98 ± 0.45, saline 1.65 ± 0.00) post administration for males (Figure 8A) and 30 minutes (194: 2.01 ± 0.37, saline 1.65 ± 0.00), 1 hour (194: 2.15 ± 0.34, saline 1.65 ± 0.00) and 2 hours (194: 2.15 ± 0.34, saline 1.65 ± 0.00) post administration in female rats (Figure 8B). This reduction demonstrates that intranasal delivery of **194** reduces mechanical allodynia induced by ION-CCI.

#### 3.7.3 Effects of intranasal administration of compound **194** on pinprick response in rats with ION-CCI

Treatment with intranasal **194** resulted in a significant decrease in the pinprick response score in the ipsilateral side of the face in rats exposed to ION-CCI compared to those treated with saline. The decrease in pinprick score was observed as early as 30 minutes post administration (**194**: 1.60 ± 0.84, saline 3.00 ± 0.47), and the effect remained significant at 1 hour (**194**: 1.7 ± 0.82, saline 2.90 ± 0.57), 2 hours (**194**: 1.80 ± 0. 79, saline 2.60 ± 0.52) and 3 hours (**194**: 1.80 ± 0. 91, saline 2.90 ± 0.32) post administration for males (Figure 8C) and 30 minutes (**194**: 1.70 ± 0.26, saline 2.90 ± 0.11) and 1 hour post administration for females (**194**: 2.00 ± 0.30, saline 3.00 ± 0.00) (Figure 8D). These findings indicate that compound **194** effectively reduces acute nociception in male and female rats with ION-CCI.

The above results highlight that intranasal administration is an effective route for delivering compound **194** to target cranial neuropathic pain.

## 4. Discussion

Despite the incidence of PTNP [5; 7], much of pain research has focused on the study of spinal nerves, and as a result, the management of craniofacial pain remains poor. In this study, we use molecular, electrophysiological, and behavioral approaches to demonstrate that regulation of Na_V_1.7 via blocking SUMOylation of CRMP2 by the small molecule **194** can be used as a therapeutic target for the alleviation of trigeminal neuropathic pain.

Although increased functionality of Na_V_1.7 is a known contributor to chronic pain, efforts to directly target Na_V_1.7 as a therapeutic have failed [24; 47; 69]. To circumvent these failures, we have advanced a differentiated strategy to allosterically regulate Na_V_1.7 [14; 15; 20; 22; 26], resulting in the discovery of compound **194** [13]. Compound **194** is a benzoylpiperidylbenzimidazole that blocks CRMP2 SUMOylation in DRG neurons and promotes internalization of Na_V_1.7, thereby decreasing sodium currents and was shown to reduce pain-like behaviors across multiple pain models, including formalin, postsurgical, spared nerve injury, spared nerve ligation, oxaliplatin- and paclitaxel-induced peripheral neuropathy, and HIV-induced sensory neuropathy in two species (mice and rats) via four routes of administration (intrathecal, intraperitoneal, subcutaneous, and oral) [13]. The efficay of **194** was not accompanied by any tolerance, addiction, rewarding properties, or neurotoxicity [13]. Here, we evaluated whether **194**’s mechanism of regulating Na_V_1.7 extended to the trigeminal ganglion. Although the presence and activity of both CRMP2 and Na_V_1.7 channels have been reported in TG neurons [54; 81; 82], this work is the first to demonstrate direct functional and molecular interactions between these two proteins in TG.

The mechanism of analgesia of **194** reportedly involves a central component and is predicted to have a high likelihood of passively diffusing through the blood-brain barrier (BBB) [13]. Studies have demonstrated that intranasal delivery can rapidly and efficiently transport drugs from the nasal cavity to the brain. This noninvasive method follows the olfactory and trigeminal nerve pathways, effectively circumventing the blood-brain barrier. Experiments conducted on rats revealed that when lidocaine is administered intranasally, structures innervated by the trigeminal nerve accumulate significantly higher concentrations of lidocaine, up to 20 times greater than those found in the brain and bloodstream. This phenomenon allows targeted therapeutic treatment for orofacial disorders, minimizing unwanted side effects in the brain and the rest of the body [37]. Furthermore, intranasal lidocaine has demonstrated analgesic effects in patients suffering from second-division trigeminal neuralgia, without causing serious adverse reactions [39]. As **194** reduced trigeminal neuropathic pain-like mechanical allodynia and acute nociception demonstrates that intranasal administration is an effective and safe route of administration for **194** to target cranial neuropathic pain.

We found that the genes for Na_V_1.7 (*Scn9a*), CRMP2 (*Dpysl2*), and Ubc9 (*Ube2i*) co-localize moslty in trigeminal A-∂ and peptidergic C fibers, which are responsible for the transmission of noxious stimuli, and our PLA experiments suggest a physical interaction between these two proteins. Approximately 75% of *Scn9a* neurons express both *Ube2i* and *Dpysl2*, which is supported by our PLA results demonstrating a physical interaction between Na_V_1.7 and CRMP2 as a result of CRMP2 SUMOylation. A more detailed characterization of this *Scn9a/Dpysl2/Ube2i* subpopulation could further direct the development of more specifically targeted drugs. Intranasal **194** was also able to effectively reduce behavioral signs of PTNP in both male and female rats in the CCI-ION model. Interestingly, we found approximately 25% of *Scn9a* cells that do not express *Ube2i*, suggesting that some *Scn9a/Dpysl2* TG cells do not undergo Ubc9-mediated SUMOylation. Therefore, the ability of **194** to reduce the behavioral signs of ION-CCI suggests that targeting a subpopulation of Na_V_1.7 is sufficient for reduction of PTNP.

We also demonstrate that over half of *Ube2i* is expressed in cells that are *Scn9a* and/or *Dpysl2* negative. This illustrates that Ubc9 SUMOylates proteins other than CRMP2 in the TG, which may include transcription factors, ion channels, and signaling proteins [2]. While many peripheral sensory neurons in the DRG undergo SUMOylation and as a result have altered functionality, the role of SUMOylation in the TG is largely understudied. Although TG and DRG are homologs of each other, differences in their transcriptional profiles have been reported at baseline [41] as well as following corresponding peripheral nerve injuries [42] suggesting that TG and DRG may respond differently to nerve injury. Therefore, aside from CRMP2, Ubc9 may contribute to SUMOylation of other molecular targets in the TG that ultimately contribute to the manifestation of craniofacial pain.

Mechanistically, we have previously shown that inhibition of CRMP2 SUMOylation triggers the assembly of an endocytic protein complex around Na_V_1.7, facilitating its internalization through a clathrin-mediated pathway ultimately leading to reduced surface expression and current density of Na_V_1.7 [33; 52]. Similarly, **194** engages this mechanism to reduce Na_V_1.7 currents by promoting clathrin-mediated internalization of Na_V_1.7 through blocking SUMOylation of CRMP2 in DRGs [13] as well as in TGs as shown in this study. Bridging the gap from molecular to organismal length scales, here, using total internal reflection fluorescence (TIRF) combined with super resolution microscopy (SIM), we report that the reduction in surface expression of Na_V_1.7 caused by **194** is due to decreased diffusivity of channels, observed at single molecule resolution for the first time, as well as an increased clustering of the immobile particles. Based on the observations of single Na_V_1.7 channels, we can infer that **194** inhibits rapid diffusion of the receptor on the membrane, which could in turn be related to channel activity as suggested by work from Akin and colleagues [1]. This remains an avenue for further investigation.

We used electrophysiology to evaluate the functional implications of Na_V_1.7/CRMP2 uncoupling, which revealed that Na_V_1.7 accounts for aproximately 50% of the total sodium currents in TG, similar to what has been previously reported in DRGs (57%) [13]. Based on the prominent involvement of Na_V_1.7 in TGs and the selectivity of **194** for Na_V_1.7 over other Na_V_ channels [13], we assessed the effect of **194** on TG neurons. Overnight treatment with **194** significantly decreased the total sodium currents of TG neurons and did not show further inhibition when coapplied along with Pro-TxII, demonstrating that **194** can modulate the entirety of Na_V_1.7 current in TG neurons. This is similar to the effect of **194** in DRG neurons but opposite to what has been found in nodose ganglia neurons, where CRMP2 does not regulate Nav1.7 and **194** has no effect [13; 46]. These conflicting findings point to a cell-specific mechanism of regulation of Na_V_1.7 channels by CRMP2 SUMOylation.

The initiation of the action potentials is due to the amplification of subthreshold depolarizations provided by Na_V_1.7 channels in sensory neurons [77]. The role of Na_V_1.7 in contributing to neuronal excitability via increased action potential firing is manifested in the human gain-of-function mutations of these channels [18]. We found that inhibition of trigeminal Na_V_1.7 via **194** decreased the number of evoked action potentials in TG neurons, suggesting that the inhibition of Na_V_1.7 via suppression of CRMP2 SUMOylation dampens the excitability of trigeminal neurons. Comparable results were observed in lumbar dorsal horn neurons, where **194** significantly decreased the number of evoked action potentials [13]. Together, the present results support the conclusion that **194** prevents the Ubc9-CRMP2 interaction and represents an indirect path to regulate Na_V_1.7 channels in TG sensory neurons.

Overall, our work demonstrates that **194** is an effective inhibitor of Na_V_1.7/CRMP2 interaction, a mechanism that is conserved among different nociceptive cellular types, with efficacy in the treatment of neuropathic pain not only spinal but also of trigeminal origin. Intranasal formulation of **194** may be a palusible route to improve treatment of trigeminal neuralgia.

## Acknowledgments

This study was supported by the NIH awards from NINDS (NS098772 and NS120663 to R.K.) and NIDA (DA042852 to R.K.). H.N.A. was funded by a postdoctoral fellowship (F32NS128392) from NINDS.

## Author Contributions

R.K. developed the concept and designed the experiments. R.K. supervised experiments and advised on data analysis. S.L.L., H.A., and R.K. wrote the manuscript. S.L.L., K.G., and A.C.-R., C.T. performed the electrophysiological recordings. P.D. performed biochemistry experiments. H.N.A performed RNAscope® fluorescence *in situ* hybridization. U.K. and R.S. carried out the behavioral experiments. S.S., R.Z. and A.D. carried out single molecule imaging and analyses. O.A.K. designed and supervised the behavioral testing studies, performed surgeries and behavioral testing. U.K. and R.S. assisted with surgeries and behavioral testing. All authors had the opportunity to discuss the results and comment on the manuscript.

## References

[1] Akin EJ, Higerd-Rusli GP, Mis MA, Tanaka BS, Adi T, Liu S, Dib-Hajj FB, Waxman SG, Dib-Hajj SD. Building sensory axons: Delivery and distribution of Na(V)1.7 channels and effects of inflammatory mediators. Sci Adv 2019;5(10):eaax4755.

[2] Anderson DB, Wilkinson KA, Henley JM. Protein SUMOylation in neuropathological conditions. Drug News Perspect 2009;22(5):255–265.

[3] Barron RP, Benoliel R, Zeltser R, Eliav E, Nahlieli O, Gracely RH. Effect of dexamethasone and dipyrone on lingual and inferior alveolar nerve hypersensitivity following third molar extractions: preliminary report. J Orofac Pain 2004;18(1):62–68.

[4] Bellampalli SS, Ji Y, Moutal A, Cai S, Wijeratne EMK, Gandini MA, Yu J, Chefdeville A, Dorame A, Chew LA, Madura CL, Luo S, Molnar G, Khanna M, Streicher JM, Zamponi GW, Gunatilaka AAL, Khanna R. Betulinic acid, derived from the desert lavender Hyptis emoryi, attenuates paclitaxel-, HIV-, and nerve injury-associated peripheral sensory neuropathy via block of N-and T-type calcium channels. Pain 2019;160(1):117–135.

[5] Benoliel R, Teich S, Eliav E. Painful Traumatic Trigeminal Neuropathy. Oral Maxillofac Surg Clin North Am 2016;28(3):371–380.

[6] Benoliel R, Wilensky A, Tal M, Eliav E. Application of a pro-inflammatory agent to the orbital portion of the rat infraorbital nerve induces changes indicative of ongoing trigeminal pain. Pain 2002;99(3):567–578.

[7] Benoliel R, Zadik Y, Eliav E, Sharav Y. Peripheral painful traumatic trigeminal neuropathy: clinical features in 91 cases and proposal of novel diagnostic criteria. J Orofac Pain 2012;26(1):49–58.

[8] Bharadwaj VN, Tzabazis AZ, Klukinov M, Manering NA, Yeomans DC. Intranasal Administration for Pain: Oxytocin and Other Polypeptides. Pharmaceutics 2021;13(7).

[9] Braden K, Stratton HJ, Salvemini D, Khanna R. Small molecule targeting NaV1.7 via inhibition of the CRMP2-Ubc9 interaction reduces and prevents pain chronification in a mouse model of oxaliplatin-induced neuropathic pain. Neurobiol Pain 2022;11:100082.

[10] Brittain JM, Duarte DB, Wilson SM, Zhu W, Ballard C, Johnson PL, Liu N, Xiong W, Ripsch MS, Wang Y, Fehrenbacher JC, Fitz SD, Khanna M, Park CK, Schmutzler BS, Cheon BM, Due MR, Brustovetsky T, Ashpole NM, Hudmon A, Meroueh SO, Hingtgen CM, Brustovetsky N, Ji RR, Hurley JH, Jin X, Shekhar A, Xu XM, Oxford GS, Vasko MR, White FA, Khanna R. Suppression of inflammatory and neuropathic pain by uncoupling CRMP-2 from the presynaptic Ca²⁺ channel complex. Nat Med 2011;17(7):822–829.

[11] Brittain JM, Piekarz AD, Wang Y, Kondo T, Cummins TR, Khanna R. An atypical role for collapsin response mediator protein 2 (CRMP-2) in neurotransmitter release via interaction with presynaptic voltage-gated calcium channels. Journal of biological chemistry 2009;284:31375–31390.

[12] Brustovetsky T, Khanna R, Brustovetsky N. CRMP2 Is Involved in Regulation of Mitochondrial Morphology and Motility in Neurons. Cells 2021;10(10).

[13] Cai S, Moutal A, Yu J, Chew LA, Isensee J, Chawla R, Gomez K, Luo S, Zhou Y, Chefdeville A, Madura C, Perez-Miller S, Bellampalli SS, Dorame A, Scott DD, Francois-Moutal L, Shan Z, Woodward T, Gokhale V, Hohmann AG, Vanderah TW, Patek M, Khanna M, Hucho T, Khanna R. Selective targeting of NaV1.7 via inhibition of the CRMP2-Ubc9 interaction reduces pain in rodents. Sci Transl Med 2021;13(619):eabh1314.

[14] Chew LA, Bellampalli SS, Dustrude ET, Khanna R. Mining the Na(v)1.7 interactome: Opportunities for chronic pain therapeutics. Biochem Pharmacol 2019;163:9–20.

[15] Chew LA, Khanna R. CRMP2 and voltage-gated ion channels: potential roles in neuropathic pain. Neuronal Signal 2018;2(1).

[16] Chi XX, Schmutzler BS, Brittain JM, Wang Y, Hingtgen CM, Nicol GD, Khanna R. Regulation of N-type voltage-gated calcium channels (Cav2.2) and transmitter release by collapsin response mediator protein-2 (CRMP-2) in sensory neurons. J Cell Sci 2009;122(Pt 23):4351–4362.

[17] Dhuria SV, Hanson LR, Frey WH, 2nd. Intranasal delivery to the central nervous system: mechanisms and experimental considerations. J Pharm Sci 2010;99(4):1654–1673.

[18] Dib-Hajj SD, Yang Y, Black JA, Waxman SG. The Na(V)1.7 sodium channel: from molecule to man. Nat Rev Neurosci 2013;14(1):49–62.

[19] Dong W, Jin SC, Allocco A, Zeng X, Sheth AH, Panchagnula S, Castonguay A, Lorenzo LE, Islam B, Brindle G, Bachand K, Hu J, Sularz A, Gaillard J, Choi J, Dunbar A, Nelson-Williams C, Kiziltug E, Furey CG, Conine S, Duy PQ, Kundishora AJ, Loring E, Li B, Lu Q, Zhou G, Liu W, Li X, Sierant MC, Mane S, Castaldi C, Lopez-Giraldez F, Knight JR, Sekula RF, Jr., Simard JM, Eskandar EN, Gottschalk C, Moliterno J, Gunel M, Gerrard JL, Dib-Hajj S, Waxman SG, Barker FG2nd, Alper SL, Chahine M, Haider S, De Koninck Y, Lifton RP, Kahle KT. Exome Sequencing Implicates Impaired GABA Signaling and Neuronal Ion Transport in Trigeminal Neuralgia. iScience 2020;23(10):101552.

[20] Dustrude ET, Moutal A, Yang X, Wang Y, Khanna M, Khanna R. Hierarchical CRMP2 posttranslational modifications control NaV1.7 function. Proc Natl Acad Sci U S A 2016;113(52):E8443–E8452.

[21] Dustrude ET, Perez-Miller S, FranÃ§ois-Moutal L, Moutal A, Khanna M, Khanna R. A single structurally conserved SUMOylation site in CRMP2 controls NaV1.7 function. Channels (Austin, Tex) 2017;11:316–328.

[22] Dustrude ET, Wilson SM, Ju W, Xiao Y, Khanna R. CRMP2 protein SUMOylation modulates NaV1.7 channel trafficking. J Biol Chem 2013;288(34):24316–24331.

[23] Ebersberger A, Portz S, Meissner W, Schaible HG, Richter F. Effects of N-, P/Q-and L-type calcium channel blockers on nociceptive neurones of the trigeminal nucleus with input from the dura. Cephalalgia 2004;24(4):250–261.

[24] Emery EC, Luiz AP, Wood JN. Nav1.7 and other voltage-gated sodium channels as drug targets for pain relief. Expert Opin Ther Targets 2016;20(8):975–983.

[25] Ershov D, Phan MS, Pylvänäinen JW, Rigaud SU, Le Blanc L, Charles-Orszag A, Conway JRW, Laine RF, Roy NH, Bonazzi D, Duménil G, Jacquemet G, Tinevez JY. TrackMate 7: integrating state-of-the-art segmentation algorithms into tracking pipelines. Nat Methods 2022;19(7):829–832.

[26] François-Moutal L, Dustrude ET, Wang Y, Brustovetsky T, Dorame A, Ju W, Moutal A, Perez-Miller S, Brustovetsky N, Gokhale V, Khanna M, Khanna R. Inhibition of the Ubc9 E2 SUMO-conjugating enzyme-CRMP2 interaction decreases NaV1.7 currents and reverses experimental neuropathic pain. Pain 2018;159(10):2115–2127.

[27] François-Moutal L, Scott DD, Perez-Miller S, Gokhale V, Khanna M, Khanna R. Chemical shift perturbation mapping of the Ubc9-CRMP2 interface identifies a pocket in CRMP2 amenable for allosteric modulation of Nav1.7 channels. Channels (Austin) 2018;12(1):219–227.

[28] François-Moutal L, Wang Y, Moutal A, Cottier KE, Melemedjian OK, Yang X, Wang Y, Ju W, Largent-Milnes TM, Khanna M, Vanderah TW, Khanna R. A membrane-delimited N-myristoylated CRMP2 peptide aptamer inhibits CaV2.2 trafficking and reverses inflammatory and postoperative pain behaviors. Pain 2015;156(7):1247–1264.

[29] Fromm GH, Terrence CF, Chattha AS. Baclofen in the treatment of trigeminal neuralgia: double-blind study and long-term follow-up. Ann Neurol 1984;15(3):240–244.

[30] Gambeta E, Gandini MA, Souza IA, Zamponi GW. Ca V 3.2 calcium channels contribute to trigeminal neuralgia. Pain 2022;163(12):2315–2325.

[31] Gomes AC, Vasconcelos BC, de Oliveira e Silva ED, da Silva LC. Lingual nerve damage after mandibular third molar surgery: a randomized clinical trial. J Oral Maxillofac Surg 2005;63(10):1443–1446.

[32] Gomez K, Duran P, Tonello R, Allen HN, Boinon L, Calderon-Rivera A, Loya-López S, Nelson TS, Ran D, Moutal A, Bunnett NW, Khanna R. Neuropilin-1 is essential for vascular endothelial growth factor A–mediated increase of sensory neuron activity and development of pain-like behaviors. PAIN 9900:10.1097/j.pain.0000000000002970.

[33] Gomez K, Ran D, Madura CL, Moutal A, Khanna R. Non-SUMOylated CRMP2 decreases Na(V)1.7 currents via the endocytic proteins Numb, Nedd4–2 and Eps15. Mol Brain 2021;14(1):20.

[34] Gregg JM. Neuropathic complications of mandibular implant surgery: review and case presentations. Ann R Australas Coll Dent Surg 2000;15:176–180.

[35] Honegger A, Spinelli S, Cambillau C, Plückthun A. A mutation designed to alter crystal packing permits structural analysis of a tight-binding fluorescein-scFv complex. Protein Sci 2005;14(10):2537–2549.

[36] Imamura Y, Kawamoto H, Nakanishi O. Characterization of heat-hyperalgesia in an experimental trigeminal neuropathy in rats. Exp Brain Res 1997;116(1):97–103.

[37] Johnson NJ, Hanson LR, Frey WH, II. Trigeminal Pathways Deliver a Low Molecular Weight Drug from the Nose to the Brain and Orofacial Structures. Molecular Pharmaceutics 2010;7(3):884–893.

[38] Kadoi J, Takeda M, Matsumoto S. Prostaglandin E2 potentiates the excitability of small diameter trigeminal root ganglion neurons projecting onto the superficial layer of the cervical dorsal horn in rats. Exp Brain Res 2007;176(2):227–236.

[39] Kanai A, Suzuki A, Kobayashi M, Hoka S. Intranasal lidocaine 8% spray for second-division trigeminal neuralgia. Br J Anaesth 2006;97(4):559–563.

[40] Klasser GD, Kugelmann AM, Villines D, Johnson BR. The prevalence of persistent pain after nonsurgical root canal treatment. Quintessence Int 2011;42(3):259–269.

[41] Kogelman LJA, Christensen RE, Pedersen SH, Bertalan M, Hansen TF, Jansen-Olesen I, Olesen J. Whole transcriptome expression of trigeminal ganglia compared to dorsal root ganglia in Rattus Norvegicus. Neuroscience 2017;350:169–179.

[42] Korczeniewska OA, Katzmann Rider G, Gajra S, Narra V, Ramavajla V, Chang YJ, Tao Y, Soteropoulos P, Husain S, Khan J, Eliav E, Benoliel R. Differential gene expression changes in the dorsal root versus trigeminal ganglia following peripheral nerve injury in rats. Eur J Pain 2020;24(5):967–982.

[43] Lechin F, van der Dijs B, Lechin ME, Amat J, Lechin AE, Cabrera A, Gomez F, Acosta E, Arocha L, Villa S, et al. Pimozide therapy for trigeminal neuralgia. Arch Neurol 1989;46(9):960–963.

[44] Li J, Stratton HJ, Lorca SA, Grace PM, Khanna R. Small molecule targeting NaV1.7 via inhibition of the CRMP2-Ubc9 interaction reduces pain in chronic constriction injury (CCI) rats. Channels (Austin) 2022;16(1):1–8.

[45] Liu CY, Lu ZY, Li N, Yu LH, Zhao YF, Ma B. The role of large-conductance, calcium-activated potassium channels in a rat model of trigeminal neuropathic pain. Cephalalgia 2015;35(1):16–35.

[46] Loya-Lopez SI, Duran P, Ran D, Calderon-Rivera A, Gomez K, Moutal A, Khanna R. Cell specific regulation of NaV1.7 activity and trafficking in rat nodose ganglia neurons. Neurobiol Pain 2022;12:100109.

[47] McDonnell A, Collins S, Ali Z, Iavarone L, Surujbally R, Kirby S, Butt RP. Efficacy of the Nav1.7 blocker PF-05089771 in a randomised, placebo-controlled, double-blind clinical study in subjects with painful diabetic peripheral neuropathy. Pain 2018;159(8):1465–1476.

[48] Meents JE, Bressan E, Sontag S, Foerster A, Hautvast P, Rosseler C, Hampl M, Schuler H, Goetzke R, Le TKC, Kleggetveit IP, Le Cann K, Kerth C, Rush AM, Rogers M, Kohl Z, Schmelz M, Wagner W, Jorum E, Namer B, Winner B, Zenke M, Lampert A. The role of Nav1.7 in human nociceptors: insights from human induced pluripotent stem cell-derived sensory neurons of erythromelalgia patients. Pain 2019;160(6):1327–1341.

[49] Messlinger K, Russo AF. Current understanding of trigeminal ganglion structure and function in headache. Cephalalgia 2019;39(13):1661–1674.

[50] Michalet X. Mean square displacement analysis of single-particle trajectories with localization error: Brownian motion in an isotropic medium. Phys Rev E Stat Nonlin Soft Matter Phys 2010;82(4 Pt 1):041914.

[51] Moon S, Lee SJ, Kim E, Lee CY. Hypoesthesia after IAN block anesthesia with lidocaine: management of mild to moderate nerve injury. Restor Dent Endod 2012;37(4):232–235.

[52] Moutal A, Cai S, Yu J, Stratton HJ, Chefdeville A, Gomez K, Ran D, Madura CL, Boinon L, Soto M, Zhou Y, Shan Z, Chew LA, Rodgers KE, Khanna R. Studies on CRMP2 SUMOylation-deficient transgenic mice identify sex-specific Nav1.7 regulation in the pathogenesis of chronic neuropathic pain. Pain 2020;161(11):2629–2651.

[53] Moutal A, Dustrude ET, Largent-Milnes TM, Vanderah TW, Khanna M, Khanna R. Blocking CRMP2 SUMOylation reverses neuropathic pain. Mol Psychiatry 2018;23(11):2119–2121.

[54] Moutal A, Eyde N, Telemi E, Park KD, Xie JY, Dodick DW, Porreca F, Khanna R. Efficacy of (S)-Lacosamide in preclinical models of cephalic pain. Pain Rep 2016;1(1).

[55] Moutal A, Li W, Wang Y, Ju W, Luo S, Cai S, Francois-Moutal L, Perez-Miller S, Hu J, Dustrude ET, Vanderah TW, Gokhale V, Khanna M, Khanna R. Homology-guided mutational analysis reveals the functional requirements for antinociceptive specificity of collapsin response mediator protein 2-derived peptides. Br J Pharmacol 2018;175(12):2244–2260.

[56] Moutal A, Luo S, Largent-Milnes TM, Vanderah TW, Khanna R. Cdk5-mediated CRMP2 phosphorylation is necessary and sufficient for peripheral neuropathic pain. Neurobiol Pain 2019;5.

[57] Moutal A, Wang Y, Yang X, Ji Y, Luo S, Dorame A, Bellampalli SS, Chew LA, Cai S, Dustrude ET, Keener JE, Marty MT, Vanderah TW, Khanna R. Dissecting the role of the CRMP2-neurofibromin complex on pain behaviors. Pain 2017;158(11):2203–2221.

[58] Moutal A, Yang X, Li W, Gilbraith KB, Luo S, Cai S, François-Moutal L, Chew LA, Yeon SK, Bellampalli SS, Qu C, Xie JY, Ibrahim MM, Khanna M, Park KD, Porreca F, Khanna R. CRISPR/Cas9 editing of Nf1 gene identifies CRMP2 as a therapeutic target in neurofibromatosis type 1-related pain that is reversed by (S)-Lacosamide. Pain 2017;158(12):2301–2319.

[59] Mustafa ER, Gambeta E, Stringer RN, Souza IA, Zamponi GW, Weiss N. Electrophysiological and computational analysis of Ca(v)3.2 channel variants associated with familial trigeminal neuralgia. Mol Brain 2022;15(1):91.

[60] Nixdorf DR, Moana-Filho EJ, Law AS, McGuire LA, Hodges JS, John MT. Frequency of persistent tooth pain after root canal therapy: a systematic review and meta-analysis. J Endod 2010;36(2):224–230.

[61] Ovesný M, Křížek P, Borkovec J, Svindrych Z, Hagen GM. ThunderSTORM: a comprehensive ImageJ plug-in for PALM and STORM data analysis and super-resolution imaging. Bioinformatics 2014;30(16):2389–2390.

[62] Piekarz AD, Due MR, Khanna M, Wang B, Ripsch MS, Wang R, Meroueh SO, Vasko MR, White FA, Khanna R. CRMP-2 peptide mediated decrease of high and low voltage-activated calcium channels, attenuation of nociceptor excitability, and anti-nociception in a model of AIDS therapy-induced painful peripheral neuropathy. Mol Pain 2012;8:54.

[63] Pigg M, Svensson P, Drangsholt M, List T. Seven-year follow-up of patients diagnosed with atypical odontalgia: a prospective study. J Orofac Pain 2013;27(2):151–164.

[64] Queral-Godoy E, Figueiredo R, Valmaseda-Castellon E, Berini-Aytes L, Gay-Escoda C. Frequency and evolution of lingual nerve lesions following lower third molar extraction. J Oral Maxillofac Surg 2006;64(3):402–407.

[65] Renton T, Adey-Viscuso D, Meechan JG, Yilmaz Z. Trigeminal nerve injuries in relation to the local anaesthesia in mandibular injections. Br Dent J 2010;209(9):E15.

[66] Saurabh S, Zhang M, Mann VR, Costello AM, Bruchez MP. Kinetically Tunable Photostability of Fluorogen-Activating Peptide-Fluorogen Complexes. Chemphyschem 2015;16(14):2974–2980.

[67] Schmalhofer WA, Calhoun J, Burrows R, Bailey T, Kohler MG, Weinglass AB, Kaczorowski GJ, Garcia ML, Koltzenburg M, Priest BT. ProTx-II, a selective inhibitor of NaV1.7 sodium channels, blocks action potential propagation in nociceptors. Mol Pharmacol 2008;74(5):1476–1484.

[68] Shields KG, Storer RJ, Akerman S, Goadsby PJ. Calcium channels modulate nociceptive transmission in the trigeminal nucleus of the cat. Neuroscience 2005;135(1):203–212.

[69] Siebenga P, van Amerongen G, Hay JL, McDonnell A, Gorman D, Butt R, Groeneveld GJ. Lack of Detection of the Analgesic Properties of PF-05089771, a Selective Na(v) 1.7 Inhibitor, Using a Battery of Pain Models in Healthy Subjects. Clin Transl Sci 2020;13(2):318–324.

[70] Smith JG, Elias LA, Yilmaz Z, Barker S, Shah K, Shah S, Renton T. The psychosocial and affective burden of posttraumatic neuropathy following injuries to the trigeminal nerve. J Orofac Pain 2013;27(4):293–303.

[71] Smith MH, Lung KE. Nerve injuries after dental injection: a review of the literature. J Can Dent Assoc 2006;72(6):559–564.

[72] Szent-Gyorgyi C, Schmidt BF, Creeger Y, Fisher GW, Zakel KL, Adler S, Fitzpatrick JA, Woolford CA, Yan Q, Vasilev KV, Berget PB, Bruchez MP, Jarvik JW, Waggoner A. Fluorogen-activating single-chain antibodies for imaging cell surface proteins. Nat Biotechnol 2008;26(2):235–240.

[73] Takeda M, Ikeda M, Takahashi M, Kanazawa T, Nasu M, Matsumoto S. Suppression of ATP-induced excitability in rat small-diameter trigeminal ganglion neurons by activation of GABAB receptor. Brain Res Bull 2013;98:155–162.

[74] Takeda M, Takahashi M, Kitagawa J, Kanazawa T, Nasu M, Matsumoto S. Brain-derived neurotrophic factor enhances the excitability of small-diameter trigeminal ganglion neurons projecting to the trigeminal nucleus interpolaris/caudalis transition zone following masseter muscle inflammation. Mol Pain 2013;9:49.

[75] Valmaseda-Castellon E, Berini-Aytes L, Gay-Escoda C. Inferior alveolar nerve damage after lower third molar surgical extraction: a prospective study of 1117 surgical extractions. Oral Surg Oral Med Oral Pathol Oral Radiol Endod 2001;92(4):377–383.

[76] Vos BP, Strassman AM, Maciewicz RJ. Behavioral evidence of trigeminal neuropathic pain following chronic constriction injury to the rat’s infraorbital nerve. J Neurosci 1994;14(5 Pt 1):2708-2723.

[77] Waxman SG, Zamponi GW. Regulating excitability of peripheral afferents: emerging ion channel targets. Nat Neurosci 2014;17(2):153–163.

[78] Wilson SM, Xiong W, Wang Y, Ping X, Head JD, Brittain JM, Gagare PD, Ramachandran PV, Jin X, Khanna R. Prevention of posttraumatic axon sprouting by blocking collapsin response mediator protein 2-mediated neurite outgrowth and tubulin polymerization. Neuroscience 2012;210:451–466.

[79] Young RF, Stevens R. Unmyelinated axons in the trigeminal motor root of human and cat. J Comp Neurol 1979;183(1):205–214.

[80] Zakrzewska JM, Chaudhry Z, Nurmikko TJ, Patton DW, Mullens LE. Lamotrigine (lamictal) in refractory trigeminal neuralgia: results from a double-blind placebo controlled crossover trial. Pain 1997;73(2):223–230.

[81] Zhang P, Gan YH. Prostaglandin E(2) Upregulated Trigeminal Ganglionic Sodium Channel 1.7 Involving Temporomandibular Joint Inflammatory Pain in Rats. Inflammation 2017;40(3):1102–1109.

[82] Zhang XY, Wu X, Zhang P, Gan YH. Prolonged PGE(2) treatment increased TTX-sensitive but not TTX-resistant sodium current in trigeminal ganglionic neurons. Neuropharmacology 2022;215:109156.

